# Scalable and Adaptive Spatiotemporal Modeling for Task-Based fMRI Analysis

**DOI:** 10.64898/2025.12.04.692389

**Authors:** Jungin Choi, Abhirup Datta, Martin A. Lindquist

## Abstract

Task-based fMRI is commonly analyzed using voxel-wise general linear models, a non-spatial scalable approach that can yield fragmented activation maps. Spatial alternatives such as kernel smoothing and Bayesian models address this but either blur activation boundaries or are computationally prohibitive at modern spatial resolutions. We introduce SPLASH (Spline-Based Processing for Localized Adaptive Spatial Hemodynamics), a spatially adaptive and scalable framework based on localized thin-plate spline regression within brain parcels. Its spatial flexibility allows SPLASH to adapt to heterogeneous cortical organization and to generalize across diverse spatial domains. Using its hierarchical structure, we introduce a two-stage selective inference procedure that ensures valid false discovery rate control at the parcel and voxel levels. In simulations, SPLASH consistently delivered the best overall performance: its MSE was typically only 20–40% of that of prior spatial models, and both FPR and FNR remained well controlled. SPLASH also remained stable across smoothing choices and required only 2% of the computation time of Bayesian spatial approaches. Applied to Human Connectome Project data, SPLASH produced sharper activation patterns consistent with the motor homunculus and demonstrated higher reproducibility. SPLASH provides a generalizable, spatially adaptive, and scalable framework that strengthens statistical inference and improves neuroscientific interpretability in large-scale fMRI studies.

## 1 Introduction

Functional Magnetic Resonance Imaging (fMRI) measures blood-oxygen-level–dependent (BOLD) fluctuations as an indirect marker of neural activity (Lindquist, 2008). In task-based studies, participants perform controlled tasks while whole-brain images are repeatedly acquired at tens of thousands of spatial locations, and neural responses are modeled through the hemodynamic response function (HRF), which links stimuli to changes in the BOLD signal. Statistical analyses aim to identify brain regions with systematic task-related activation, requiring models that capture both HRF-driven temporal dynamics and spatial dependencies (Worsley et al., 2004).

### 1.1 The Classical GLM Framework

The general linear model (GLM) has been the standard framework for task-based fMRI for nearly three decades (Friston et al., 1994). For an experiment with *K* task conditions, the observed BOLD signal **y** at a given spatial location is modeled as the sum of task-related responses, each obtained by convolving a stimulus function **s**_*k*_ with its HRF **h**_*k*_, plus temporal noise ***ϵ***:

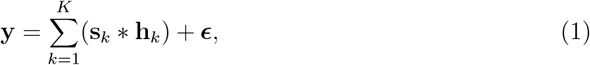

where ∗ denotes temporal convolution. Expanding the convolution,

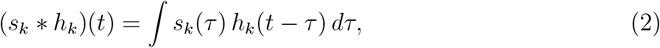

shows how each stimulus *s*_*k*_ is temporally filtered by the HRF.

Because HRF shapes naturally vary across brain regions and individuals (Lindquist et al., 2007; Lindquist and Wager, 2008), each task-specific HRF is often represented using a flexible temporal basis expansion:

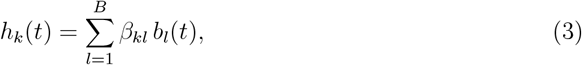

where 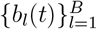 are basis functions (e.g., canonical HRF (Friston et al., 1998), B-splines (Degras and Lindquist, 2014), or FIR (Dale, 1999)), and *β*_*kl*_ are location-specific coefficients. Substituting the basis expansion into (1) yields

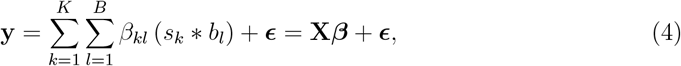

where **y** ∈ ℝ^*T*^ is the time series of length *T*, and the design matrix **X** ∈ ℝ^*T×*(*KB*)^ is constructed from the convolved basis regressors,

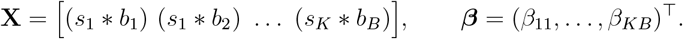

We assume ***ϵ*** ~ 𝒩 (**0, Σ**). In practice, **X** is often augmented with nuisance regressors (e.g., motion and low-frequency drift), but we omit these for clarity.

This formulation reduces inference to linear regression, enabling closed-form estimation, scalable computation, and flexible HRF parameterization. However, fitting the GLM independently at each location ignores spatial correlations, often yielding noisy, fragmented activation maps and inflated false positives (Lindquist and Mejia, 2015), particularly in high-resolution datasets such as the HCP (Van Essen et al., 2013).

### 1.2 Gaussian Kernel Smoothing

A widely used approach to incorporate spatial structure into task-based fMRI analysis is to apply Gaussian kernel smoothing (GKS) to the BOLD signal prior to GLM fitting (Worsley et al., 2004). Let the observed BOLD signal at location *v* be denoted *y*^*v*^, and denote the smoothed version by

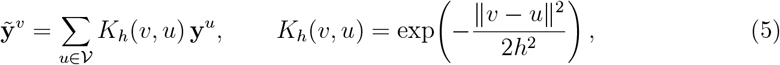

where *K*_*h*_(*v, u*) is a Gaussian kernel and *h* is the spatial bandwidth. In practice, the kernel weights are typically normalized so that they sum to one within a local neighborhood of *v*. GKS increases the signal-to-noise ratio (SNR) by aggregating information from neighboring locations and can reduce misregistration across subjects. It performs spatial smoothing directly on the data, independently of the GLM structure and the activation coefficients that are ultimately of inferential interest. Its key limitation is the use of a fixed global bandwidth (Yue et al., 2010). Fine focal activations become overly blurred, while more diffuse signals remain under-smoothed, leading to an inflated spatial extent of activation and reduced anatomical specificity. In addition, GKS applies smoothing across the entire brain and does not respect cortical folding or tissue boundaries, allowing signal to leak across sulci and mix activity from anatomically distinct regions (Glasser et al., 2016). These limitations motivate spatially adaptive methods that smooth at the level of activation estimates, in a way that better aligns with local signal structure and brain geometry.

### 1.3 Spatial Bayesian Modeling

Bayesian approaches incorporate spatial dependence in task-based fMRI by placing probabilistic priors on the activation coefficients, typically modeling spatially smooth coefficient maps rather than smoothing the raw BOLD data. Over the past two decades, Bayesian spatial modeling has evolved from early smoothing priors and Markov random field (MRF) formulations to increasingly flexible and scalable frameworks; see Zhang et al. (2015) for an overview. Early work used spatial autoregressive or MRF-type priors to enforce local smoothness across neighboring locations (Descombes et al., 1998; Woolrich et al., 2004), while later models combined these ideas with mixture structures and Dirichlet process clustering to capture latent spatial heterogeneity and jointly infer activation clusters (Hartvig and Jensen, 2000; Zhang et al., 2016). On the cortical surface, Mejia et al. (2020) introduced a surface-based Bayesian GLM using SPDE-based Gaussian Markov random field (GMRF) priors with the Integrated Nested Laplace Approximation (INLA) (Rue et al., 2009) for scalable uncertainty quantification.

Despite these advances, spatial Bayesian models remain computationally demanding for whole-brain fMRI analyses, where fully Bayesian inference via Markov chain Monte Carlo (MCMC) quickly becomes infeasible when modeling spatial dependence across the entire brain. To improve scalability, variational inference methods have been explored as approximate Bayesian alternatives (Penny et al., 2005; Zhang et al., 2016). SPDE–INLA approaches require solving GMRFs on triangulated meshes (Lindgren et al., 2011; Rue et al., 2009), which is scalable on two-dimensional cortical surfaces but becomes infeasible for three-dimensional volumetric data due to the size of the required tetrahedral meshes and associated precision matrices (Mejia et al., 2020). Fixed or globally pooled regularization parameters may oversmooth focal activations or undersmooth more diffuse patterns. Thus, although Bayesian methods operate naturally on the model coefficients and offer a principled framework for spatial regularization, their substantial computational cost and reliance on global smoothing parameters motivate the need for more spatially adaptive and computationally tractable alternatives.

### 1.4 A New Approach

We introduce SPLASH (Spline-Based Processing for Localized Adaptive Spatial Hemodynamics), a spatially adaptive framework for task-based fMRI that augments the classical GLM with localized spline-based modeling and hierarchical error control. Within predefined functional parcels, SPLASH models activation coefficients as smooth spatial functions using thin-plate regression splines, enabling smoothing in coefficient space rather than on the raw BOLD signal. Because spline bases are constructed separately within parcels, smoothing adapts to local geometry and functional heterogeneity, preserving sharp boundaries while denoising homogeneous regions. The coordinate-based thin-plate formulation also makes SPLASH applicable to both volumetric and cortical surface data.

For group-level inference, SPLASH uses its hierarchical structure in a two-stage selective testing procedure: parcels are first screened for activation using aggregated statistics, and location-specific tests are then performed only within selected parcels with adjustments that control the average false discovery rate across selected families. This yields activation maps that are sharper and more spatially coherent than those from fixed-kernel smoothing, while remaining computationally efficient and scalable than fully Bayesian spatial models.

The remainder of the paper is organized as follows. Section 2 presents the SPLASH methodology, including localized spline construction and hierarchical selective inference. Section 3 evaluates performance through simulation studies, Section 4 demonstrates SPLASH on Human Connectome Project motor-task data, and Section 5 concludes with implications and directions for future research.

## 2 Proposed Spatial Framework: SPLASH

Figure 1 summarizes the overall SPLASH workflow. SPLASH estimates activation coefficients at each spatial location (voxels or vertices) within a set of predefined parcels using localized thin plate spline regression, enabling spatially adaptive smoothing. These localized estimates are then passed to a two-level hierarchical inference procedure that provides statistically rigorous identification of activated regions through selective FDR control.

**Figure 1.**
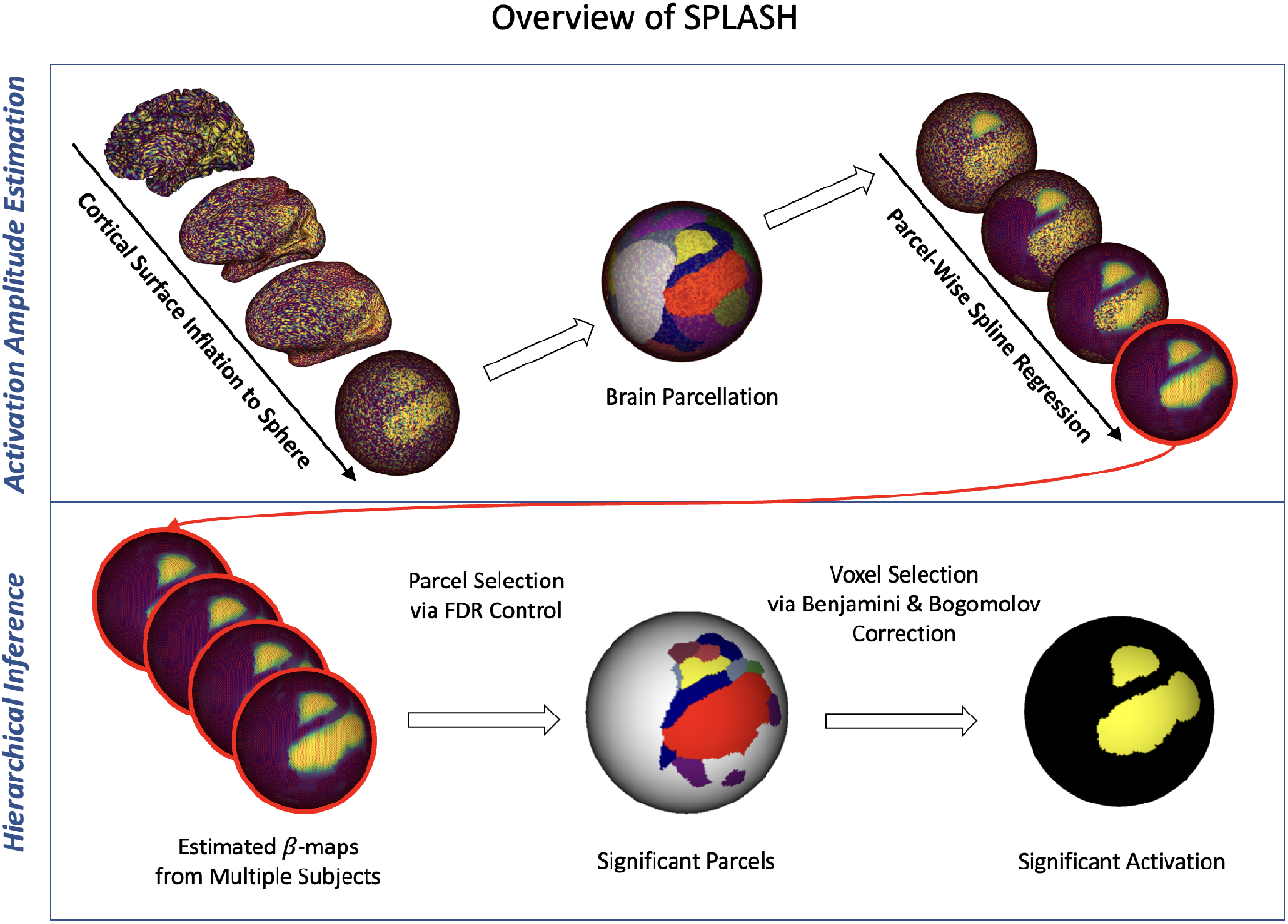
Overview of the SPLASH workflow. The upper panel illustrates localized spline-based modeling of task-related activation amplitudes. The lower panel depicts the hierarchical selective inference procedure used to identify significant activation.

### 2.1 Model Specification

Brain locations are commonly grouped into functional “parcels” with covarying activity at rest or during tasks, providing a structured representation of large-scale brain organization. Standard parcellations include the Yeo–Buckner atlas (Yeo et al., 2011), the Schaefer atlas (Schaefer et al., 2018), and other resting-state–based maps such as those of Power, Glasser, and Shen (Power et al., 2011; Glasser et al., 2016; Shen et al., 2013).

SPLASH operates independently within each parcel, enabling region-specific spatial adaptivity. Within a parcel, spatial variation in activation coefficients is modeled using thin-plate spline regression (Wood, 2003), which flexibly captures local structure while avoiding over-smoothing across anatomical boundaries. Splines are constructed using the mgcv package in R (Wood, 2017), and the spline basis dimension is selected automatically according to the parcel’s spatial heterogeneity.

For each parcel, SPLASH models spatial variation in the activation coefficients for each task and temporal basis function. Let 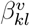 denote the coefficient for task *k* and basis *l* at location *v* from (4). For fixed *k* and *l*, the vector of coefficients across the *V* locations in a parcel is

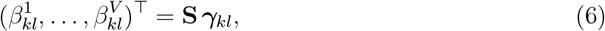

where **S** is the *V × M* thin-plate spline design matrix constructed from spatial coordinates, and ***γ***_*kl*_ is the corresponding spline coefficient vector.

This formulation separates temporal and spatial structure: temporal variation is captured by the *B* HRF basis functions and spatial variation by the *M* spline basis functions. Representing activation fields via (6) allows SPLASH to estimate the lower-dimensional spline coefficients ***γ***_*kl*_ rather than the full set 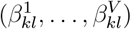, yielding both regularization and spatial adaptivity.

Having expressed the location-specific activation coefficients using the spatial spline expansion (6), the parcel-wise model can be written compactly as

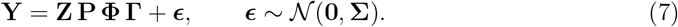

Here

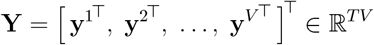

is the stacked vector of BOLD time series across the *V* locations in the parcel and

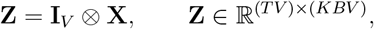

where **I**_*V*_ is the *V × V* identity matrix and **X** ∈ ℝ^*T×KB*^ is the GLM design matrix defined in (4). Thus, **Z** is block diagonal, with each block equal to **X**. The permutation matrix

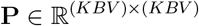

reorders the coefficient vector from location-major ordering 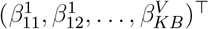 to basis-major ordering 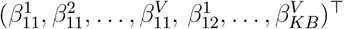, ensuring compatibility with the spline representation. The spatial basis matrix is

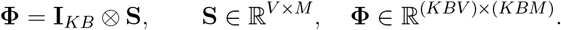

This yields the coefficient vector

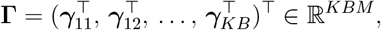

with each ***γ***_*kl*_ containing the *M* spline coefficients for task *k* and basis function *l*.

The residual vector ***ϵ*** follows a multivariate normal distribution with block-diagonal co-variance **Σ**, where each location’s temporal noise follows an AR(*p*) process, allowing location-specific temporal autocorrelation while remaining computationally tractable.

### 2.2 Model Fitting

Because the block-diagonal noise covariance **Σ** is unknown, SPLASH uses a feasible generalized least squares (FGLS) estimator (Amemiya, 1985), replacing **Σ** with an estimate 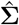 obtained from location-specific AR(*p*) models:

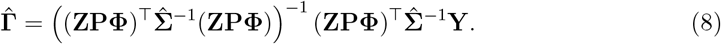

For each location, the AR(*p*) temporal covariance block is Toeplitz, and both the covariance and its inverse admit fast *O*(*T* ^2^) algorithms via the Levinson–Durbin recursion (Brockwell and Davis, 1991). Moreover, under an AR(*p*) model, the inverse covariance is banded with bandwidth *p* (Pourahmadi, 1999), making the inversion of each *T × T* block computationally efficient even when *T* is moderately large (e.g., *T* ≈ 600). As a result, the FGLS estimator remains a computationally tractable, closed-form estimator for the SPLASH model.

For independent errors, the estimator reduces to ordinary least squares (OLS):

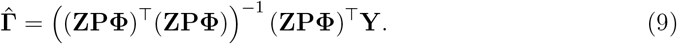

Finally, the estimated location-specific activation coefficients are reconstructed by converting back from the spline-basis representation:

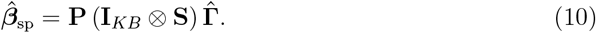

For each parcel, the number of spline basis functions *M* is selected using the Bayesian Information Criterion (BIC) (Schwarz, 1978), carefully balancing model fit and complexity. This data-driven choice provides greater flexibility in geometrically complex regions and stronger regularization in smoother areas, yielding an anatomically informed and scalable, spatially adaptive activation model.

### 2.3 Theoretical Properties

SPLASH provides several theoretical guarantees that distinguish it from GKS. Because smoothing occurs in parameter space rather than on the observed BOLD signal, SPLASH preserves the GLM estimand under mild regularity conditions and avoids distortions introduced by smoothing the raw time series. Key theoretical results are summarized below, with full derivations in the Supplementary Material.

#### 2.3.1 Bias of GKS

Unlike SPLASH, GKS smooths the observed BOLD signal before GLM fitting, producing systematic bias in both the estimated activation coefficients and their variances. The following results formalize these effects.

##### Proposition 1

(Bias of GKS activation estimates).

*Fix a parcel and the GLM in Equation* (4) *at each location v* ∈ 𝒱 *with design matrix* **X** ∈ ℝ^*T×KB*^ *(full column rank), activation vector* ***β***^*v*^ ∈ ℝ^*KB*^, *and noise* ***ϵ***^*v*^ *satisfying* 𝔼 (***ϵ***^*v*^) = **0** *and* Cov(***ϵ***^*v*^) = **Σ**_*ϵ*_.

*Let GKS be defined as in Equation* (5), *producing smoothed time series* 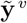 *with kernel weights summing to one, and define the estimator* 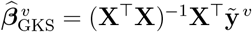.

*Then, conditional on* {***β***^*u*^}_*u*∈*V*_ *and* **X**,

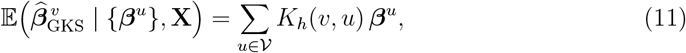

*and the resulting bias is*

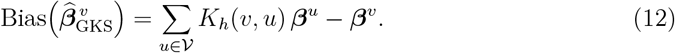

*Thus, the estimator is unbiased only when* ***β***^*u*^ *is constant over the kernel support; otherwise, GKS induces spatial smoothing bias*.

##### Proposition 2

(Variance attenuation under GKS-based GLM).

*Under the assumptions of Proposition 1, further assume temporal errors are independent across locations with common covariance* **Σ**_*ϵ*_. *Let* **R** = **I**_*T*_ − **X**(**X**^⊤^**X**)^−1^**X**^⊤^ *be the residual-forming matrix and define the GKS and unsmoothed residuals* 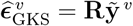 *and* 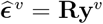.

*Then*

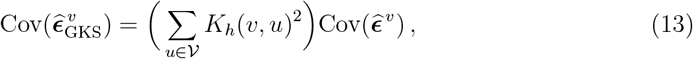

*where* 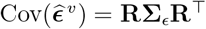.

*If the kernel is nontrivial (i*.*e*., *some u*_1_≠ *u*_2_ *satisfy K*_*h*_(*v, u*_*j*_) *>* 0*), then* 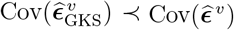 *in strict Loewner order*.

##### Corollary 1

(Variance underestimation under homoskedastic errors).

*Under the setting of Proposition 2, suppose further that temporal errors are homoskedastic and uncorrelated in time, with* 𝔼 (***ϵ***^*u*^) = **0** *and* Cov(***ϵ***^*u*^) = *σ*^2^**I**_*T*_ *for all u* ∈ 𝒱. *Then*

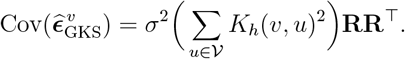

*If* 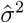 *denotes the usual unbiased estimator from the unsmoothed GLM*, 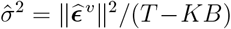 *then the analogous GKS-based estimator* 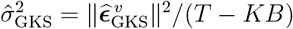 *satisfies*

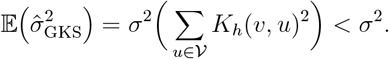

*Thus, GKS always induces downward bias in the usual variance estimator, with attenuation determined by the kernel weights*.

#### 2.3.2 Unbiasedness of SPLASH coefficients

In contrast, SPLASH solves a correctly specified linear regression in the spline basis. As a result, its coefficient estimates preserve the GLM estimand and avoid the biases of GKS.

##### Theorem 1

(Consistency of SPLASH activation estimates).

*Consider the SPLASH model in Equation* (7),

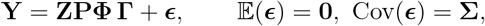

*where the design matrix* **A** = **ZPΦ** ∈ ℝ^(*TV*)*×*(*KBM*)^ *has full column rank, and* **Γ** ∈ ℝ^*KBM*^ *is the vector of spline coefficients. Let* 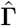 *be the FGLS estimator defined in Equation* (8), *and assume the following:*

1. *The number of independent experimental units N (e*.*g*., *subjects) tends to infinity, while the dimensions TV and KBM of* **Y** *and* **A** *remain fixed*.
2. 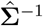 *is consistent, in the sense that*

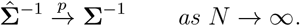
3. **A** *has full column rank, so that* **A**^⊤^**Σ**^−1^**A** *is positive definite*.

*Then the feasible GLS estimator is consistent:*

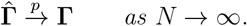

*Finally, the spatially smoothed estimates in Equation* (10) *satisfy* 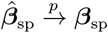.

##### Corollary 2

(Unbiasedness under independent homoskedastic errors).

*Under the SPLASH model in Equation* (7) *with fixed full-rank design matrix* **A** = **ZPΦ**, *suppose that*

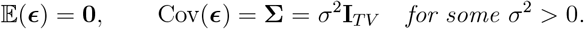

*If, in addition, the working covariance used in FGLS is restricted to the form* 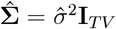 *then the FGLS estimator reduces exactly to ordinary least squares:*

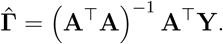

*Consequently*,

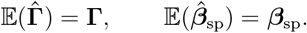

*Thus, in the independent homoskedastic setting, SPLASH with OLS yields exactly unbiased estimates of the activation field*.

#### 2.3.3 Variance estimation under SPLASH

Because SPLASH fits an exact linear model in coefficient space, its uncertainty quantification retains standard regression properties.

##### Theorem 2

(Consistency of SPLASH variance estimation).

*Under the assumptions of Theorem 1, let*

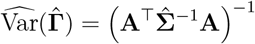

*denote the usual plug-in covariance estimator for the FGLS estimator* 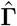.

*Then, as N* → ∞,

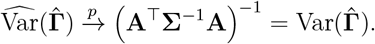

*Moreover, for any fixed linear transformation* **L** ∈ ℝ^*r×KBM*^, 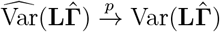.

*In particular, taking* **L** = **P**(**I**_*KB*_ ⊗ **S**) *yields*

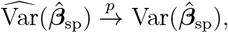

*so SPLASH provides asymptotically valid uncertainty quantification for the smoothed activation field*.

##### Corollary 3

(Unbiased variance under independent errors).

*In the setting of Corollary 2, suppose the error variance σ*^2^ *is estimated by*

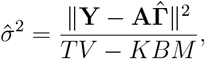

*and define*

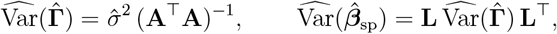

*with* **L** = **P**(**I**_*KB*_ ⊗ **S**) *as above*.

*Then*

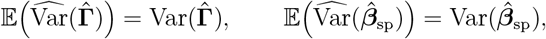

*so SPLASH achieves finite-sample unbiased variance estimation under independent errors*.

Together, these results highlight the fundamental distinction between GKS and SPLASH. GKS alters both the mean and variance of the GLM by smoothing the observed data, whereas SPLASH performs spatial smoothing in coefficient space through a correctly specified linear model, providing asymptotically unbiased activation estimates and valid variance estimation under standard GLS assumptions.

### 2.4 Hierarchical Inference with Selective Error Control

For group-level inference, we apply a two-stage hierarchical testing procedure. Throughout, we assume the data are measured over a set of voxels that make up a brain volume; all results carry over directly to surface-based analyses over cortical vertices. Parcels are first screened for activation, and voxel-wise tests are then performed only within selected parcels. To account for this selection step and control the average voxel-level false discovery rate across selected parcels, we use the selective inference adjustment of Benjamini and Bogomolov (2014) at level *α*_voxel_.

#### 2.4.1 Parcel-level screening

Let *N* be the number of participants, *P* the number of parcels, and *V*_*p*_ the number of voxels in parcel *p*. For participant *i*=1,…,*N*, let 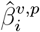 denote the estimated activation coefficient at voxel *v* in parcel *p*. Throughout this subsection we fix a task *k* and temporal basis index *l* and suppress (*k, l*) in the notation, so 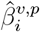 represents 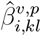. The participant-level parcel average is 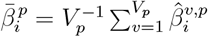, and the between-subject parcel mean is 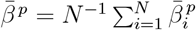.

Assuming independent observations across participants with finite variance, the parcel-level test statistic is the one-sample *t*-statistic

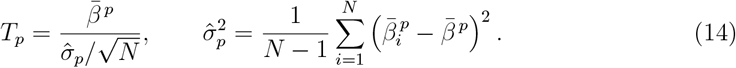

Let 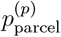 denote the two-sided *p*-value associated with *T*_*p*_. We apply the Benjamini– Hochberg procedure at level *α*_parcel_ to identify significant parcels:

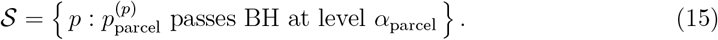

#### 2.4.2 Voxel-level inference within selected parcels

For each selected parcel *p* ∈ 𝒮, we perform voxel-level inference across participants.

The group-level mean and variance at voxel *v* are

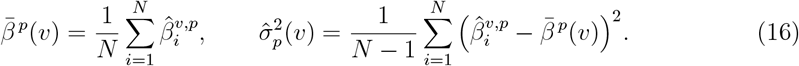

The voxel-level test statistic is the one-sample *t*-statistic

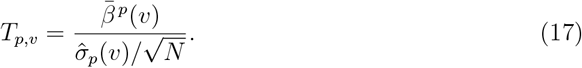

We apply the Benjamini–Hochberg (BH) procedure separately within each selected parcel *p*, using an adjusted FDR level

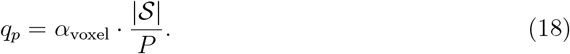

This adjustment follows the selective inference framework of Benjamini and Bogomolov (2014), which guarantees that the expected average FDR over the randomly selected parcels satisfies

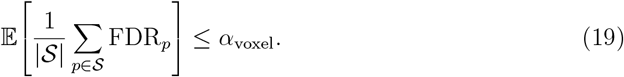

where FDR_*p*_ denotes the false discovery rate within parcel *p* (and the average is defined as 0 when |𝒮| = 0).

Thus, the two-stage procedure first identifies parcels likely to contain activation, then performs voxel-level inference only within those parcels. This selective FDR adjustment preserves power while ensuring valid error control under data-driven parcel selection.

In both Sections 3 and 4, we set *α*_parcel_ = 0.1 and *α*_voxel_ = 0.05. A slightly more liberal parcel-level threshold helps avoid missing truly active parcels, while maintaining strict voxel-level control within the selected parcels.

### 2.5 Application Domains and Spatial Representations

A key strength of SPLASH is its use of thin-plate splines, which can be constructed on any spatial domain with defined coordinates. As illustrated in Figure 2, spline bases naturally adapt to the underlying geometry, whether volumetric or surface-based, providing flexible spatial representations across diverse domains.

**Figure 2.**
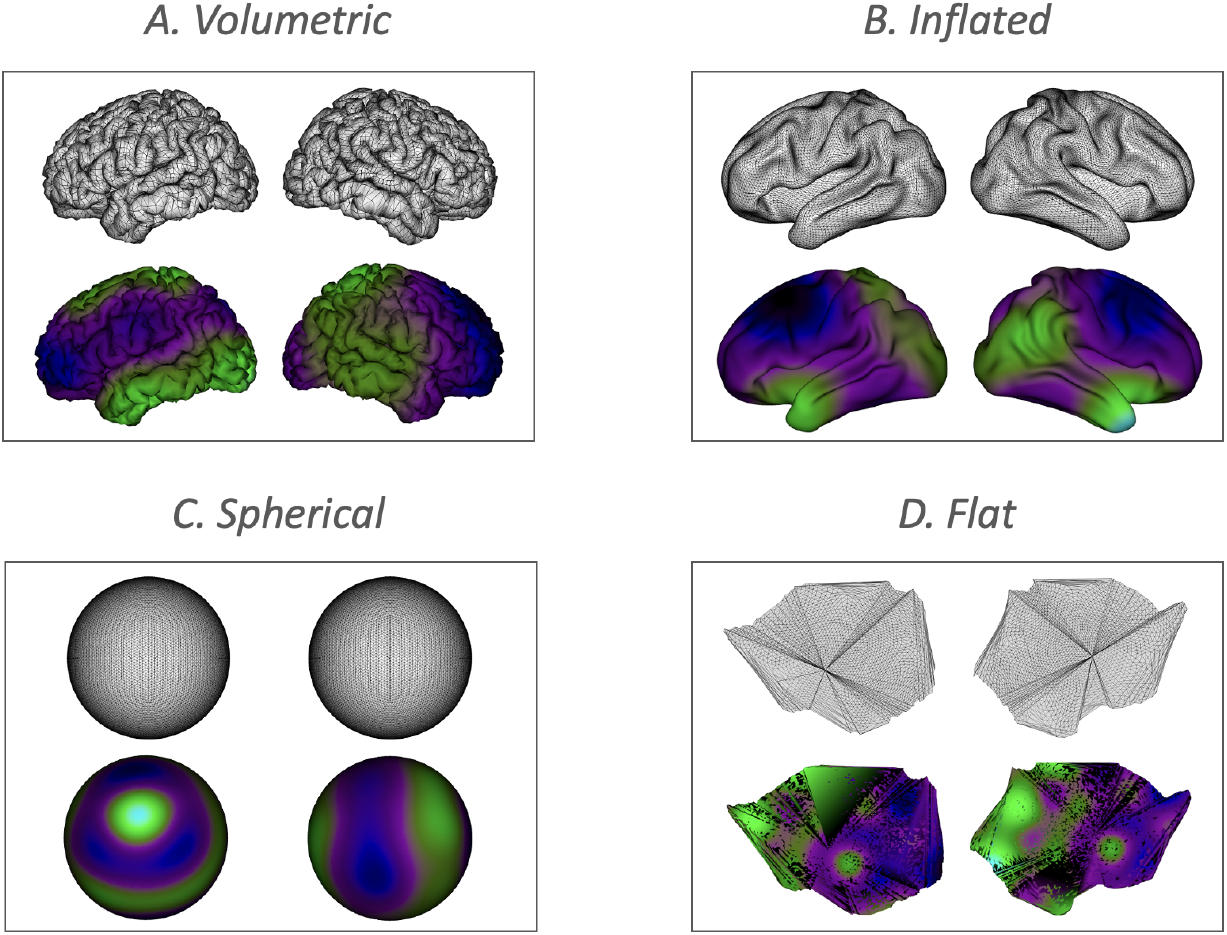
Examples of spatial domains to which SPLASH can be applied. The upper row illustrates different volumetric and surface-based geometries, and the lower row shows the corresponding thin-plate spline basis functions constructed on each domain.

In contrast, Bayesian spatial models based on SPDE–INLA (Mejia et al., 2020) are, in practice, typically applied to two-dimensional cortical surfaces, since they require a triangulated mesh and do not scale feasibly to dense three-dimensional volumetric domains due to the size and complexity of the resulting precision matrices. SPLASH avoids these limitations, offering a scalable, domain-agnostic framework that applies equally well to 3D volumes and 2D cortical surfaces without dimensional restrictions.

### 2.6 Computational Benefits

Beyond its theoretical guarantees, SPLASH is computationally efficient compared with Bayesian spatial methods such as SPDE–INLA (Mejia et al., 2020; Rue et al., 2009), which require solving large sparse precision systems on cortical meshes and do not readily extend to full volumetric domains. In contrast, SPLASH retains the linear structure of the GLM, enabling fast estimation via standard linear algebra. Smoothing parameters are selected via BIC, ensuring adaptivity without manual tuning, and the spline basis scales naturally to both volumetric (3D) and surface (2D) geometries. As a result, SPLASH provides a unified, computationally scalable, and theoretically principled framework for spatially adaptive fMRI activation modeling.

## 3 Simulation Study

### 3.1 Simulation Design

We conducted simulation to evaluate SPLASH for group-level fMRI activation mapping. Synthetic BOLD signals were generated on a spherical cortical surface with 32,492 vertices, each assigned a parcel label *p* from the Schaefer-100 parcellation (Schaefer et al., 2018).

SPLASH was compared with four methods: (i) a vertex-wise GLM without spatial smoothing, (ii) Gaussian Kernel Smoothing (GKS) with bandwidth *h* = 2.55 (equivalent to FWHM = 6), (iii) adaptive GKS (AGKS) with BIC-selected bandwidth, and (iv) a Bayesian spatial model using SPDE–INLA implemented in `bayesfMRI` (Spencer et al., 2022). Because SPLASH also adapts model complexity using BIC, AGKS and SPLASH provide parallel data-driven smoothing frameworks for comparison.

For each task *k* = 1, 2, the ground-truth activation at vertex *v* was 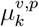 for *v* = 1, …, 32,492, where *p* is the parcel containing *v*. Each 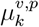 was generated as a smooth spatial field and scaled to [0, 1]. Because parcels are used only during model fitting, the true field does not vary within parcels, so 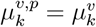.

For participant *i* = 1, …, 30, task *k*, and vertex *v*, activation coefficients were

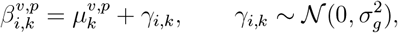

introducing between-participant variability while preserving the same spatial field. Because only one temporal basis function (canonical HRF) is used 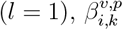 corresponds directly to 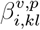 in Equation (6).

Observed time series were generated as

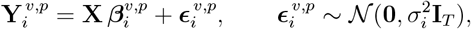

where 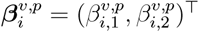, **X** is the design matrix from canonical-HRF–convolved stimuli, and *T* is the number of time points in each series. Subject-specific noise variances 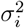 were drawn from [1, 5] to reflect realistic variability.

We examined three effective SNR levels by varying the between-participant variance:

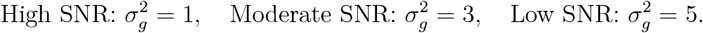

### 3.2 Results

We evaluated each method along four key dimensions: estimation accuracy, activation detection (Task 1 under the High-SNR setting), robustness to smoothing-parameter selection, and computational scalability. Additional estimation and inference results for the Moderate- and Low-SNR settings, as well as for Task 2 under High SNR, are provided in the Supplementary Material (Figures S1–S10). Numerical summaries for all five methods across all SNR levels and both tasks are reported in Tables S1–S2.

#### 3.2.1 Estimation Accuracy

Estimation performance was quantified using mean squared error (MSE) between the estimated activation maps 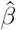 and the ground-truth field *µ*. Under high SNR, Figure 4 shows clear differences across methods. The non-spatial GLM produced fragmented and noisy estimates, reflecting its inability to borrow spatial information. GKS and AGKS reduced noise but blurred activation boundaries, particularly in the smaller activation region, and fixed-bandwidth GKS undersmoothed larger areas. The Bayesian model yielded smoother estimates with improved boundary recovery but remained less precise than SPLASH. SPLASH most accurately recovered fine activation structure while preserving boundaries, achieving the lowest MSE (0.94).

**Figure 3.**
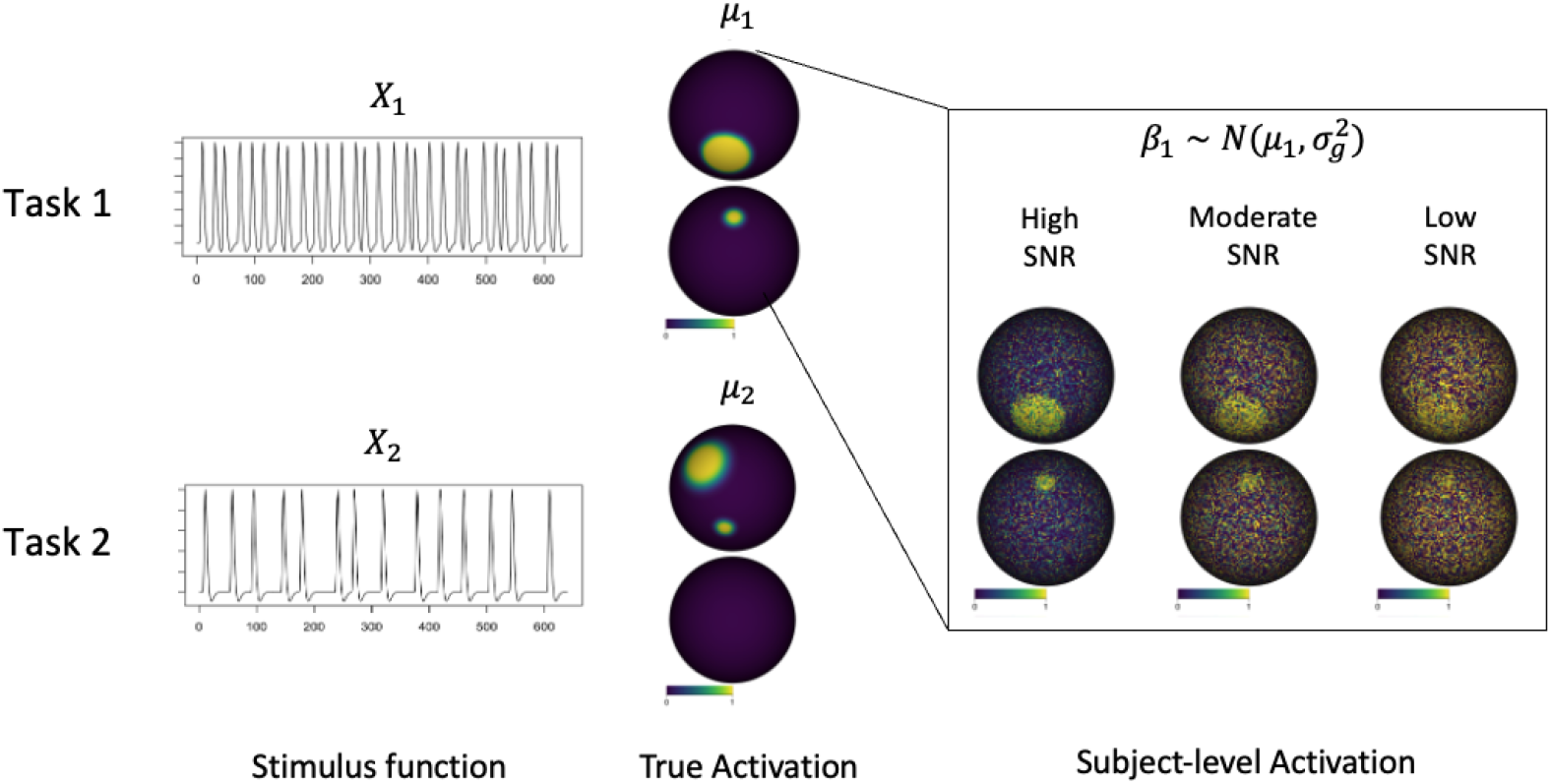
Simulation setup for generating activation coefficients and observed BOLD signals under different SNR levels.

**Figure 4.**
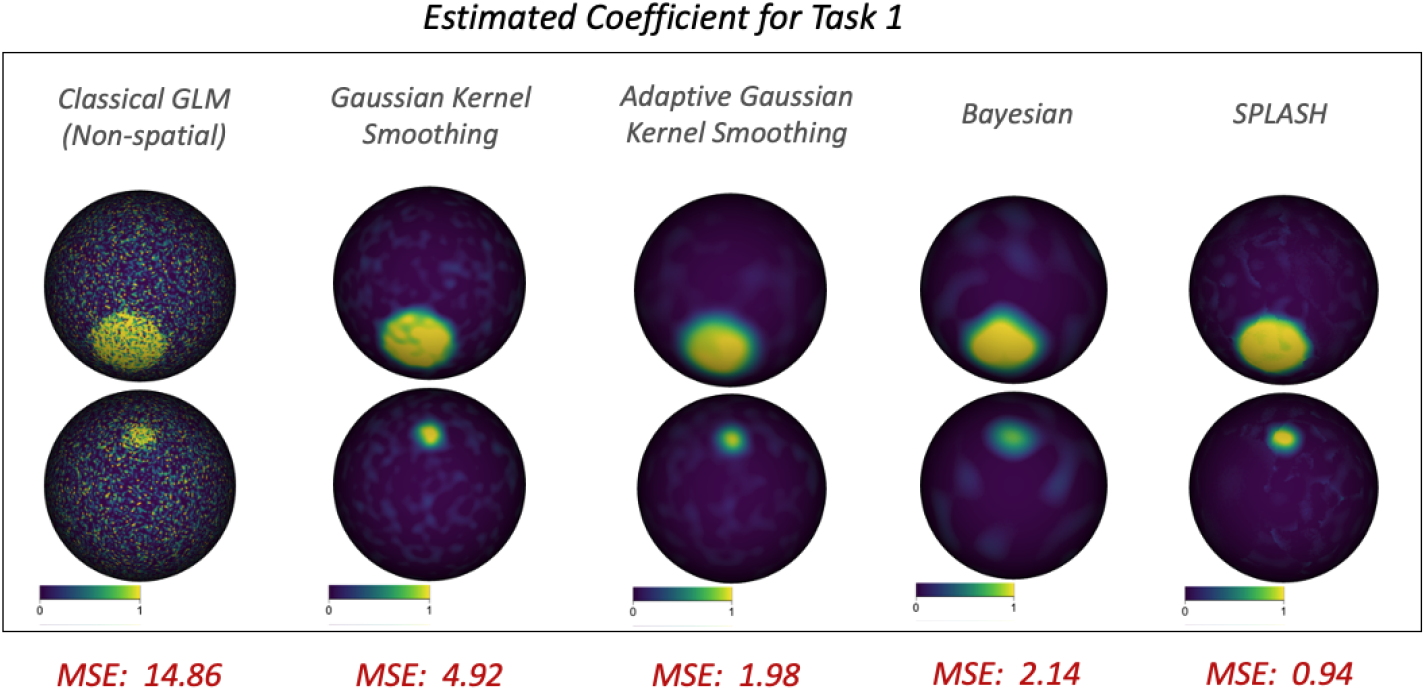
Estimated activation coefficients 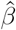 for Task 1 across methods. Bottom labels report MSE *×* 100 for each method.

#### 3.2.2 Activation detection

Activation detection for GLM, GKS, and AGKS used vertex-wise *t*-tests with Benjamini– Hochberg FDR control (Benjamini and Hochberg, 1995). The Bayesian model used posterior activation probabilities, while SPLASH employed hierarchical selective FDR control. Figure 5 shows detected activation regions, with false positive rate (FPR) and false negative rate (FNR) in red. The GLM produced scattered, salt-and-pepper detections; GKS and AGKS frequently missed localized activation due to oversmoothing. The Bayesian model better preserved spatial structure, but SPLASH achieved the lowest FNR and the second-lowest FPR, yielding the most accurate activation detection among the methods.

**Figure 5.**
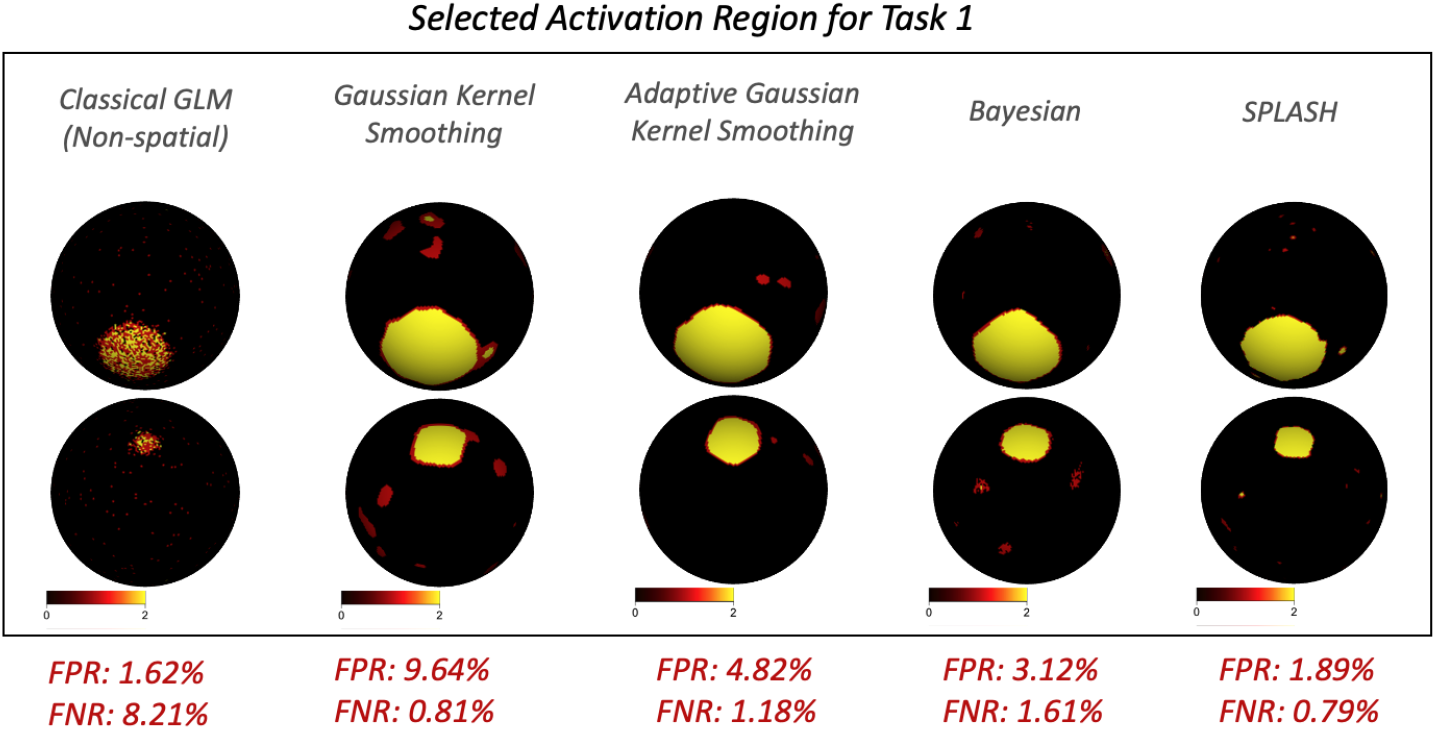
Selected activation regions for Task 1 across methods. False positive (FPR) and false negative (FNR) rates are shown in red labels below each map.

To evaluate performance across thresholds, we computed AUC values derived from ROC curves (Figure 6). SPLASH consistently achieved the highest AUC across SNR conditions for both tasks, demonstrating robust activation detection even when noise levels increased.

**Figure 6.**
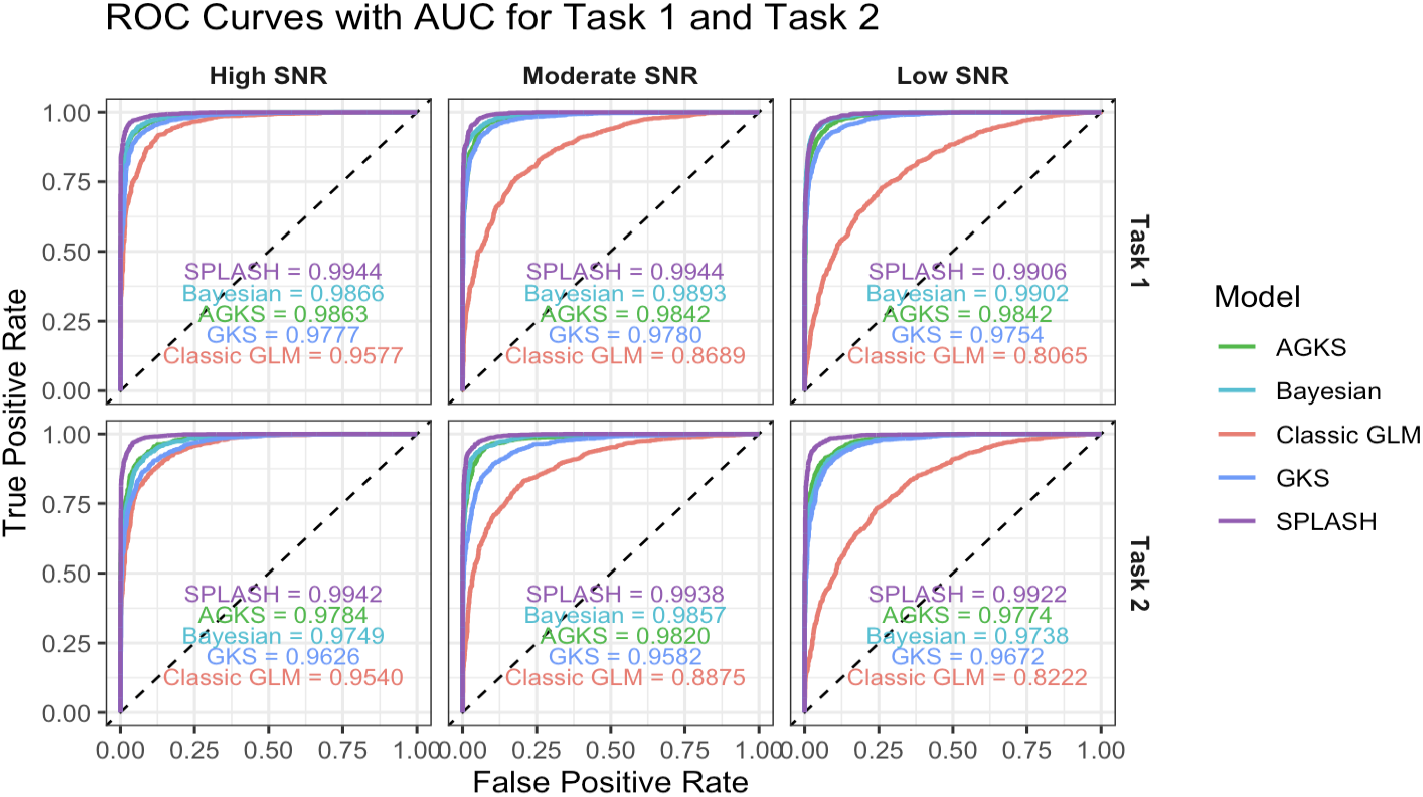
ROC curves for Task 1 and Task 2 across SNR levels. SPLASH achieves the highest AUC across all noise conditions, indicating more accurate and robust activation detection than GLM, GKS, AGKS, the Bayesian spatial model.

#### 3.2.3 Robustness to Smoothing Parameters

Both SPLASH and AGKS adapt their spatial smoothing using BIC, selecting the number of spline basis functions and the kernel bandwidth, respectively. As shown in Figure 7, GKS is highly sensitive to the choice of bandwidth *h* in Equation (5), with substantial variation in false negative and false positive rates. In contrast, SPLASH is strongly robust: varying the number of spatial spline functions *M* (the spatial basis dimension in Equation (6)) yields nearly identical activation maps and similar FNR/FPR values. This stability reflects the spatial adaptiveness of thin-plate splines, which capture heterogeneous structure without precise tuning of *M*. Additional numerical results for both methods are reported in Table S3 of the Supplementary Material.

**Figure 7.**
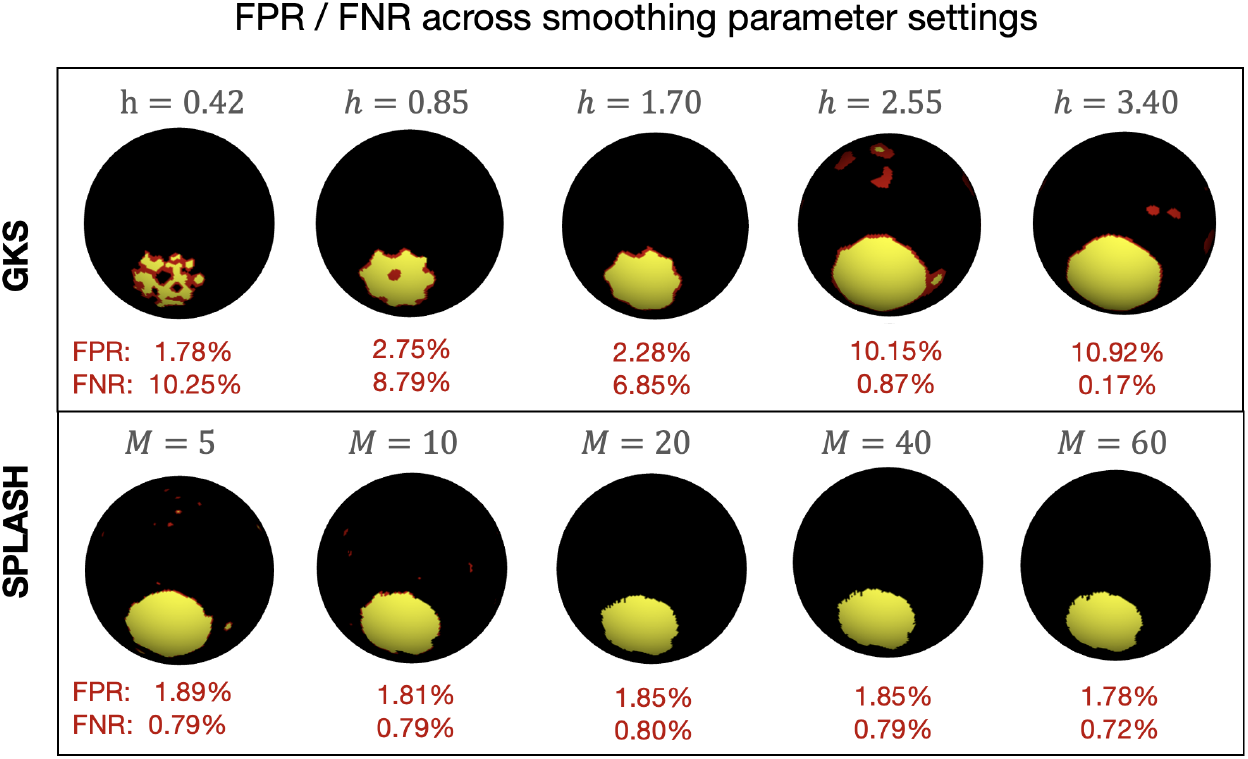
Visualization of smoothing-parameter choices under High SNR. Top: GKS results for kernel bandwidths *h*; Bottom: SPLASH results for numbers of spline basis functions *M*. Red labels show FPR and FNR averaged across Task 1 and Task 2; activation maps correspond to part of Task 1.

Estimation accuracy and activation-detection performance across smoothing parameters for GKS and SPLASH are shown in Figure 8. SPLASH exhibits stable FPR, FNR, and MSE across parameter values, indicating weak dependence on the choice of spatial basis size. In contrast, GKS is highly sensitive to the kernel bandwidth, with performance varying markedly as *h* changes, leading FPR and FNR to range from below 1% to over 10%. This robustness highlights SPLASH’s practical advantage, as it reduces the need for careful, dataset-specific tuning.

**Figure 8.**
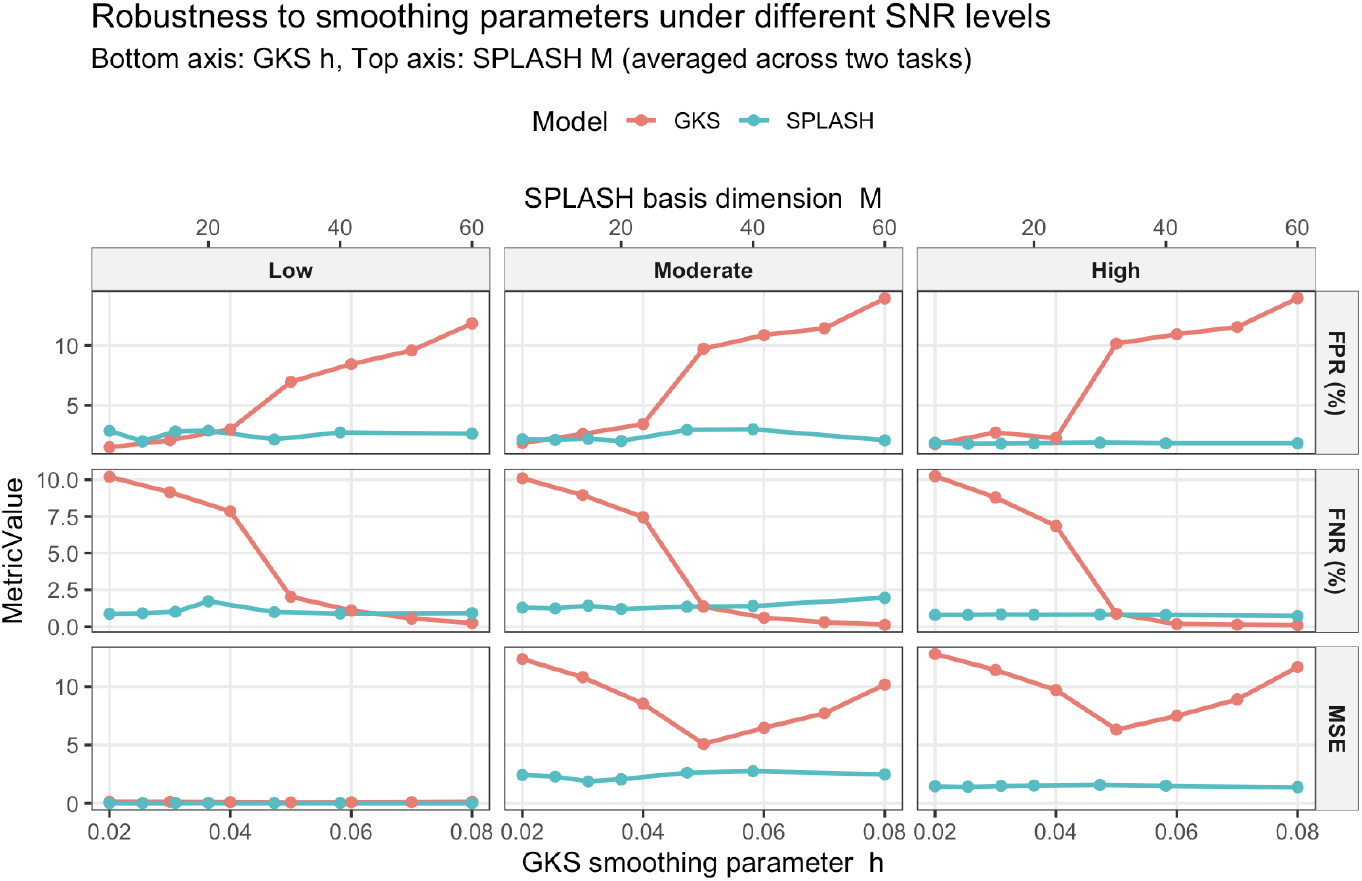
FPR (top), FNR (middle), and MSE (bottom) for GKS and SPLASH across a range of smoothing parameters under different SNR settings. Bottom axis shows the kernel bandwidth, while the top axis shows the number of spatial splines for SPLASH.

#### 3.2.4 Computational cost

For reference, we compared runtime against a spatial Bayesian model implemented in `bayesfMRI` (Spencer et al., 2022). For a single participant on the simulated cortical surface, Bayesian fitting required approximately 845 seconds, whereas SPLASH completed the analysis in roughly 7 seconds, a difference that scales substantially in group-level studies (Figure 9). All computations were run on a SLURM high-performance computing cluster (Intel Xeon Gold 6248, 40 cores, 192 GB RAM, no GPU). SPLASH was run single-threaded, while bayesfMRI used INLA’s default multithreading.

**Figure 9.**
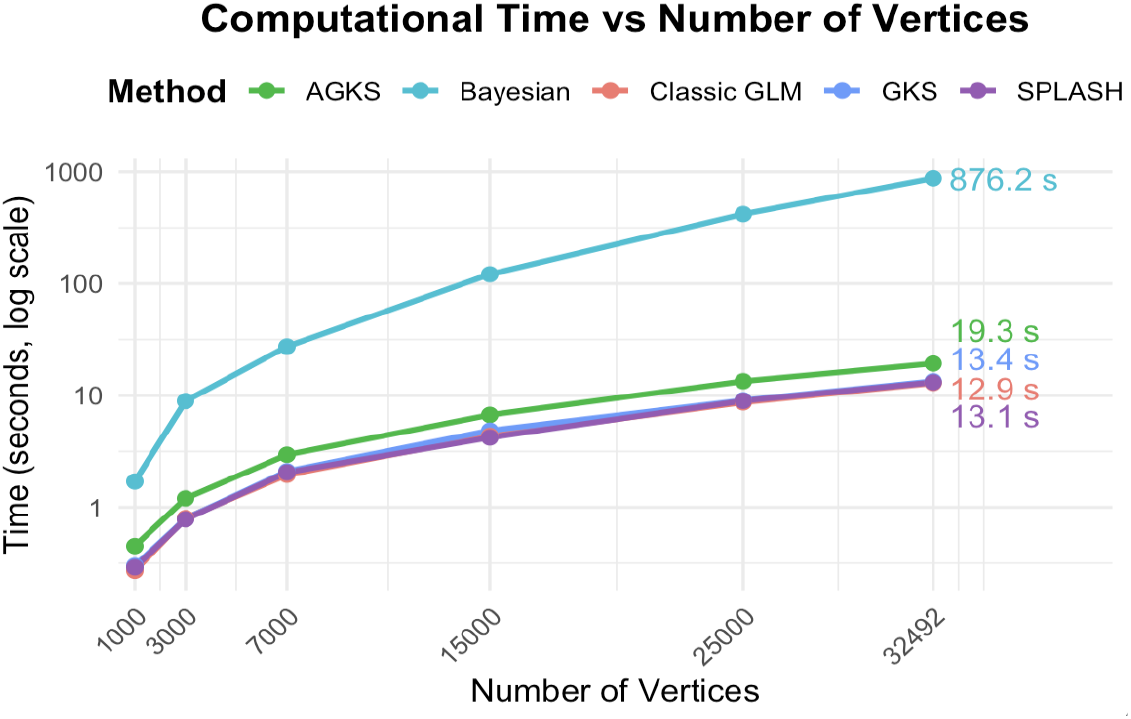
Computation time for spatial model fitting. SPLASH shows a substantial runtime advantage over Bayesian spatial modeling.

#### 3.2.5 Summary

Overall, SPLASH showed superior performance over classical GLM, GKS, AGKS, and the Bayesian spatial model. It provided lower MSE in recovering activation maps, higher detection accuracy across SNR levels, better preservation of spatial detail, and substantially improved computational scalability. These results demonstrate that SPLASH effectively balances noise suppression with spatial specificity, making it well-suited for large-scale, group-level fMRI analysis.

## 4 Application: HCP Motor Data

We applied SPLASH to motor-task cortical surface fMRI (cs-fMRI) data from *N* = 416 genetically unrelated participants in the Human Connectome Project–Young Adult (HCP-YA) dataset (Van Essen et al., 2013). Each participant completed a 3.5-minute run (*T* = 284 volumes) involving five cued movements (tongue, left/right hand, left/right foot). We analyzed minimally preprocessed cs-fMRI data mapped to the fsLR-32k surface (Fischl, 2012), including motion and distortion correction, cortical reconstruction, and spherical registration. SPLASH was fit directly on the fsLR-32k spherical surface, with fourteen nuisance regressors (six motion parameters, their derivatives, and linear/quadratic trends) removed prior to modeling to ensure cleaner, preprocessed signals.

AGKS and SPLASH were applied under three commonly used parcellations—the Yeo 17-network atlas, Schaefer-100, and Schaefer-400 (Yeo et al., 2011; Schaefer et al., 2018)—and fit independently within each parcel with smoothing selected via BIC. We also included the classical vertex-wise GLM, which was fit at each vertex without spatial smoothing. Classical GLM and AGKS used standard vertex-wise BH FDR control, whereas SPLASH employed hierarchical selective FDR consistent with its multilevel structure.

Because ground truth is unknown, performance was assessed under sample variability using 10-fold cross-validation. Participants were randomly split into ten folds (41–42 each). For each fold, models were trained on the remaining nine folds and evaluated on the held-out fold. Repeating this across folds allowed us to compare estimation accuracy and activation detection to assess robustness, generalizability, and stability under participant-level variation.

### 4.1 Results

The results in this section focus on the Tongue task using the Schaefer-100 parcellation. Subject-level and group-level analyses for the remaining tasks (Left Foot, Right Foot, Left Hand, and Right Hand), as well as SPLASH results under the other parcellations, are provided in the Supplementary Material (S11–S22).

#### 4.1.1 Subject-level Analysis

Figure 10 shows single-subject activation results for the tongue motor task. The top row displays estimated activation coefficients 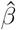 for the classical GLM, GKS, and SPLASH. The GLM yields noisy and spatially fragmented estimates, whereas GKS reduces noise but oversmooths spatial detail. SPLASH produces smoother and more spatially coherent activation patterns while retaining fine-grained structure.

**Figure 10.**
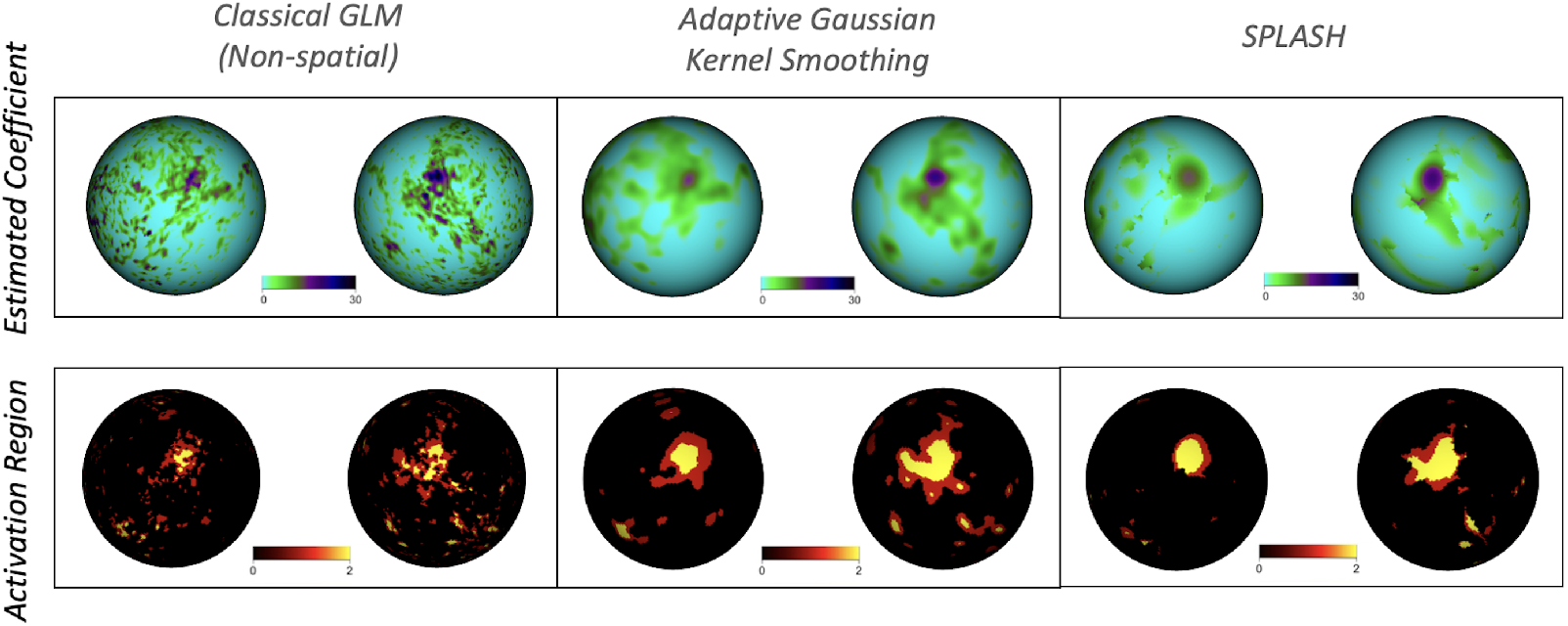
Single-subject results for the tongue motor task. Columns show classical GLM (left), AGKS (middle), and SPLASH (right). **Top row:** estimated activation coefficients 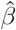. **Bottom row:** detected activation regions.

The bottom row of Figure 10 shows the corresponding detected activation regions. SPLASH identifies localized tongue activations more consistently with the motor homunculus (Penfield and Boldrey, 1937) than the GLM and with fewer missed detections than both the classical GLM and GKS, demonstrating improved sensitivity while maintaining vertex-level specificity.

#### 4.1.2 Group-level Analysis

We next examined group-level activation for the tongue motor task using all *N* = 416 participants. Across methods, the expected motor activation pattern was recovered. However, notable differences emerged in spatial specificity and noise suppression. The classical GLM produced fragmented and noisy estimates, while GKS improved spatial smoothness but remained sensitive to bandwidth choice and often obscured focal activations. SPLASH yielded the most coherent activation patterns, preserving localized motor representations while suppressing noise.

Figure 11 summarizes these results. The top row shows estimated group activation coefficients, and the bottom row displays detected activation regions for each method. SPLASH produced more spatially coherent and anatomically localized activation patterns, aligning well with the motor homunculus and yielding fewer false positives with clearer maps overall. In contrast, GKS yielded oversmoothed detections, while the classical GLM produced fragmented activation patterns lacking spatial clarity.

**Figure 11.**
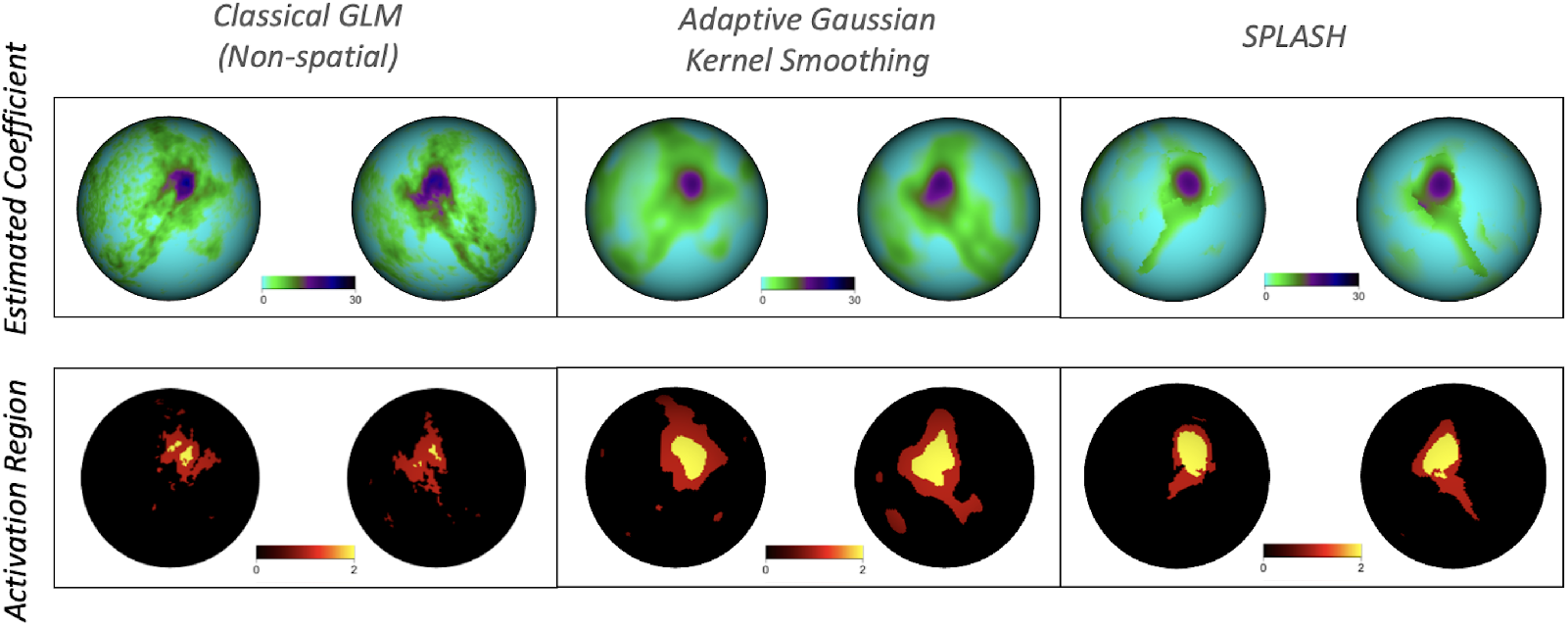
Group-level results for the tongue motor task. Columns show classical GLM (left), AGKS (middle), and SPLASH (right). **Top row:** estimated group activation coefficients. **Bottom row:** detected activation regions.

#### 4.1.3 Robustness to Parcellation

Because SPLASH is applied independently within user-defined parcels, it is important to assess whether its performance depends on the parcellation. We evaluated robustness using three widely used cortical parcellations: the Yeo 17-network atlas (Yeo et al., 2011), the Schaefer-100 atlas, and the Schaefer-400 atlas (Schaefer et al., 2018), spanning 51, 100, and 400 parcels within our analysis domain.

As shown in Figure 12, SPLASH yields highly consistent activation patterns and coefficient estimates across all parcellations, with only minor differences arising from spatial granularity. These results demonstrate that SPLASH is robust to parcellation choice and produces stable activation estimates across diverse schemes. Additional results for other motor tasks are provided in the Supplementary Material (S19–S22).

**Figure 12.**
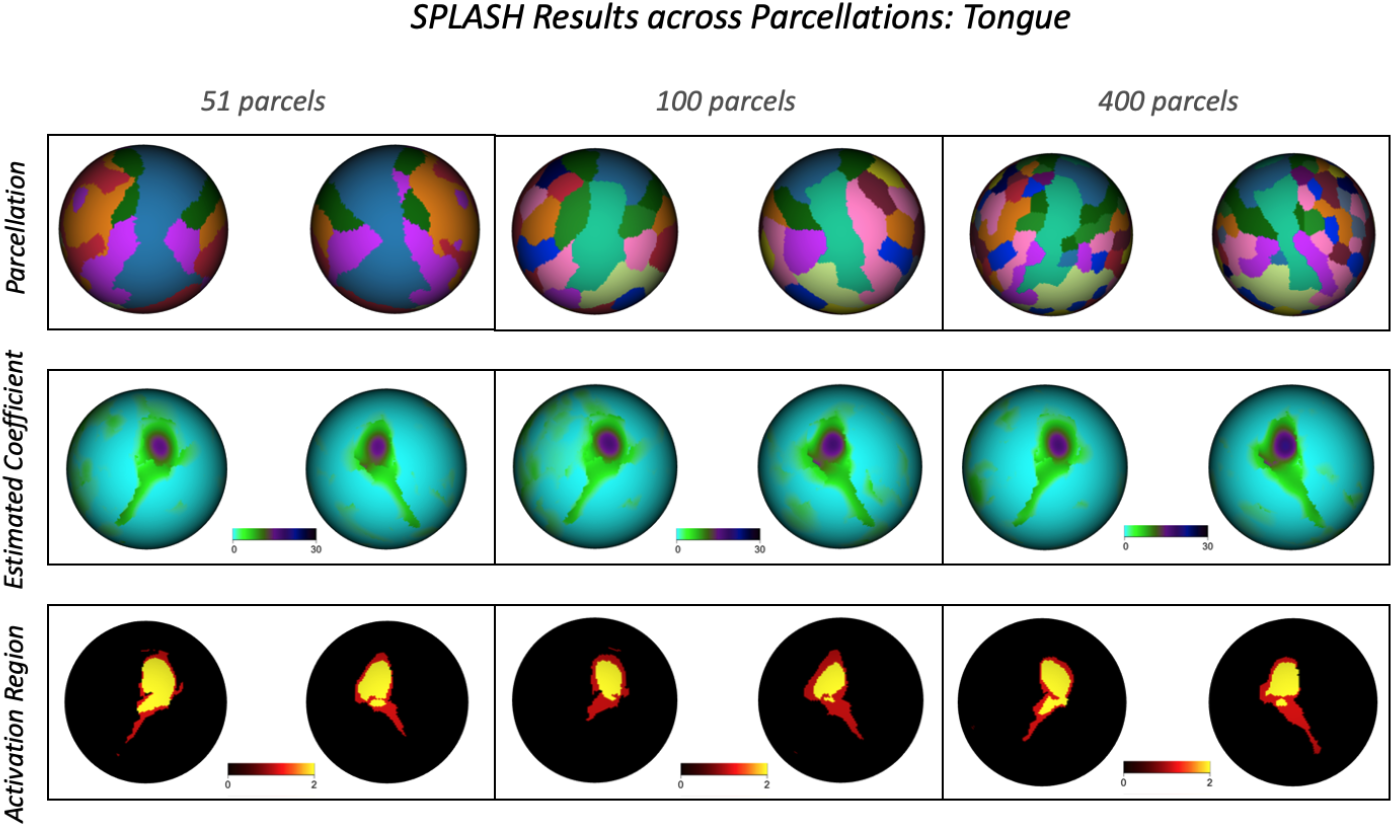
Top row: cortical parcellations used in the robustness analysis (Yeo 17, Schaefer 100, and Schaefer 400). Middle row: SPLASH-estimated activation coefficients for the tongue task under each parcellation. Bottom row: detected activation regions (yellow: *p <* 0.01, red: *p <* 0.05).

#### 4.1.4 10-fold Cross-Validation

We further assessed model accuracy, generalizability, and robustness using 10-fold cross-validation. Because real data lack ground-truth activation, we quantified reproducibility with False Positive Concordance (FPC) and False Negative Concordance (FNC), which measure how consistently each method identifies activation across folds (Figure 13). FPC and FNC denote the proportions of vertices declared active-but-inactive and inactive-but-active relative to the activation map from the other nine folds, respectively. Thus, lower FPC and FNC indicate better agreement with this cross-validated reference.

**Figure 13.**
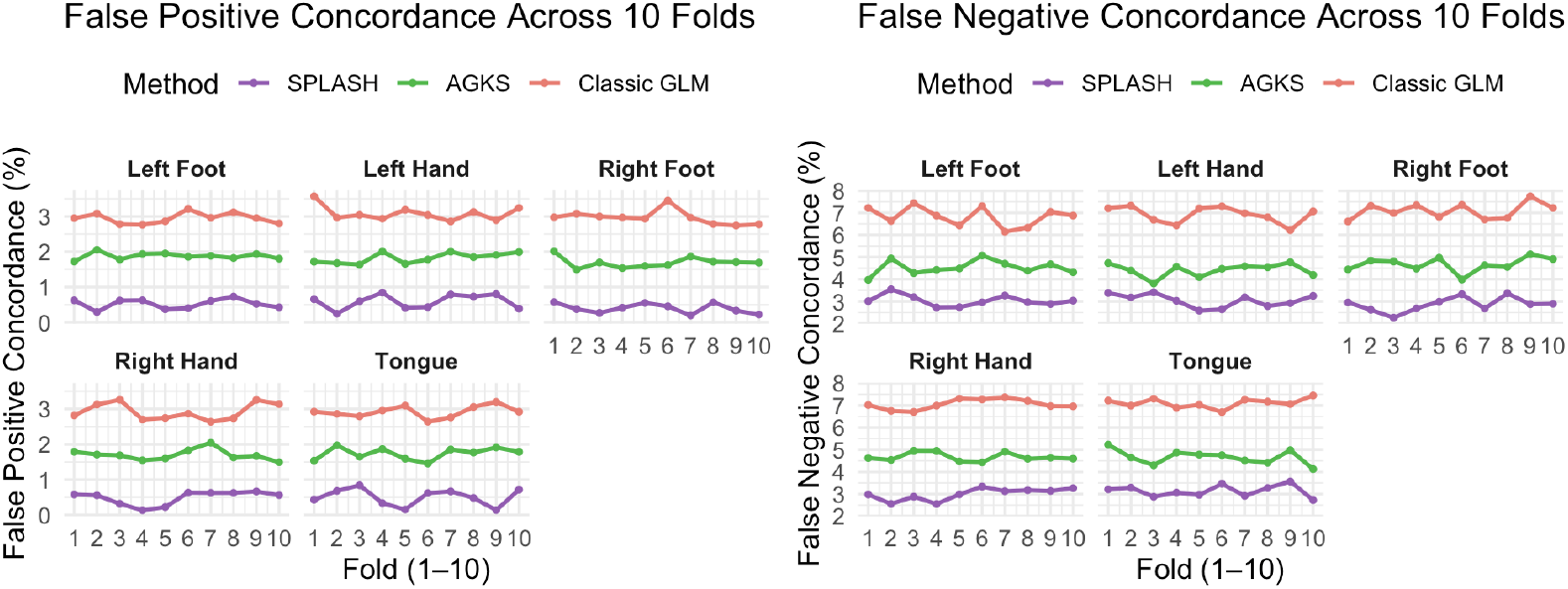
10-fold cross-validation concordance for motor task activation. Curves show FPC and FNC for each method across folds, summarizing the stability and reproducibility of activation detection.

SPLASH achieved both the lowest FPC and FNC across 10 folds, demonstrating more reproducible activation patterns across folds. These results indicate that SPLASH improves the reliability of group-level inference beyond gains in spatial estimation alone.

## 5 Discussion

SPLASH provides a spatially adaptive framework for task-based fMRI that bridges the gap between fixed-kernel smoothing and fully Bayesian spatial modeling. By fitting localized spline regressions within parcels and selecting basis complexity via BIC, SPLASH tailors smoothing to local signal structure, preserving sharp activation boundaries while reducing noise in homogeneous regions. This yields activation estimates that are both interpretable and stable across subjects and data resampling.

Across simulations and HCP data, SPLASH improved spatial estimation and activation detection over voxel-wise GLM and both fixed-bandwidth and adaptive Gaussian kernel smoothing. In particular, SPLASH mitigates the variance underestimation that can inflate false discoveries under kernel smoothing, leading to more reliable empirical FPR and FNR control and greater reproducibility in 10-fold cross-validation. The method is computationally efficient because inference reduces to standard linear algebra, enabling scalability to high-resolution surface datasets that remain challenging for spatial Bayesian methods.

Bayesian spatial models, including SPDE–INLA formulations, offer a principled approach to uncertainty quantification and continuous spatial smoothing but remain computationally intensive for large fMRI cohorts. SPLASH achieves spatial adaptivity with substantially lower computational burden, making it a practical alternative for large-scale neuroimaging studies where thousands of subjects or repeated model fits are required.

Because SPLASH operates within a predefined cortical parcellation, its performance may depend on how well the chosen parcellation aligns with true functional boundaries. Our main analysis used the Schaefer-100 atlas, and we demonstrated robustness across the Yeo 17-network and Schaefer-400 parcellations. Nonetheless, alternative or multiscale atlases may offer different levels of spatial adaptiveness or capture complementary organizational features. Future work may therefore evaluate SPLASH across a broader set of parcellation schemes or develop data-driven, adaptive parcellations that adjust to observed activity patterns, potentially improving flexibility and reliability in heterogeneous brain regions.

Although this work focuses on spatial modeling for activation detection, SPLASH naturally extends to joint spatial–temporal analysis, including nonparametric HRF estimation (Goutte et al., 2000; Lindquist et al., 2007). Integrating SPLASH with smooth FIR-based HRF modeling may improve sensitivity to regional differences in neurovascular coupling. Additional extensions include incorporating structured priors for multi-subject borrowing of strength, connecting SPLASH with hierarchical Bayesian inference, and adapting the method for naturalistic or rapid event-related designs.

In summary, SPLASH provides a statistically principled and computationally scalable approach to adaptive spatial modeling in fMRI, improving accuracy, false discovery control, and reproducibility, and making it well suited for large neuroimaging studies where spatial heterogeneity and efficiency are key considerations.

