## Supplementary Material for "Scalable and Adaptive Spatiotemporal Modeling for Task-Based fMRI Analysis"

### Supplementary Materials for: Adaptive and Scalable Spatiotemporal Modeling for Task-Based fMRI Data

### Overview

This document contains supplementary material for the manuscript. It includes mathematical proofs, expanded simulation studies, robustness analyses, and additional empirical results referenced in the main text.

### 1 Proofs of Theoretical Properties

This section provides the full statements and complete proofs of all propositions, corollaries, and theorems referenced in Section 2.3 of the main text.

**Proposition 1** (Bias of GKS activation estimates)**.** *Fix a parcel and consider the GLM*

$$\mathbf{y}^v = \mathbf{X}\boldsymbol{\beta}^v + \boldsymbol{\epsilon}^v, \qquad v \in \mathcal{V},$$

*where $\mathbf{y}^v \in \mathbb{R}^T$ is the observed BOLD time series at location $v$, $\mathbf{X} \in \mathbb{R}^{T\times(KB)}$ is a fixed full-column-rank design matrix, $\boldsymbol{\beta}^v \in \mathbb{R}^{KB}$ is the vector of activation coefficients, and $\boldsymbol{\epsilon}^v$ is zero-mean noise with*

$$\mathbb{E}(\boldsymbol{\epsilon}^v) = \mathbf{0}, \qquad \mathrm{Cov}(\boldsymbol{\epsilon}^v) = \boldsymbol{\Sigma}_\epsilon,$$

*for all $v \in \mathcal{V}$. No assumptions are made about spatial dependence across locations.*

*Let the Gaussian kernel smoothing (GKS) operator at location $v$ be*

$$\tilde{\mathbf{y}}^{\,v} = \sum_{u\in\mathcal{V}} K_h(v,u)\,\mathbf{y}^u,$$

*with weights satisfying $K_h(v,u) \geq 0$ and $\sum_u K_h(v,u) = 1$. Define the GKS-based GLM estimator*

$$\widehat{\boldsymbol{\beta}}^v_{\mathrm{GKS}} = (\mathbf{X}^\top\mathbf{X})^{-1}\mathbf{X}^\top\tilde{\mathbf{y}}^{\,v}.$$

*Then, conditional on $\{\boldsymbol{\beta}^u\}_{u\in\mathcal{V}}$ and $\mathbf{X}$,*

$$\mathbb{E}\Big(\widehat{\boldsymbol{\beta}}^v_{\mathrm{GKS}} \mid \{\boldsymbol{\beta}^u\}, \mathbf{X}\Big) = \sum_{u\in\mathcal{V}} K_h(v,u)\,\boldsymbol{\beta}^u. \tag{1}$$

*Consequently, the bias at location $v$ is*

$$\mathrm{Bias}\Big(\widehat{\boldsymbol{\beta}}^v_{\mathrm{GKS}}\Big) = \sum_{u\in\mathcal{V}} K_h(v,u)\,\boldsymbol{\beta}^u - \boldsymbol{\beta}^v, \tag{2}$$

*In particular, the bias is zero if $\boldsymbol{\beta}^u$ is constant over all $u$ such that $K_h(v,u) > 0$, and is* *generally nonzero whenever $\boldsymbol{\beta}$ varies within the kernel support.*

*Proof.* Fix a location $v \in \mathcal{V}$. By definition of the GKS operator,

$$\tilde{\mathbf{y}}^{\,v} = \sum_{u \in \mathcal{V}} K_h(v,u)\,\mathbf{y}^u.$$

Substituting the GLM $\mathbf{y}^u = \mathbf{X}\boldsymbol{\beta}^u + \boldsymbol{\epsilon}^u$ gives

$$\tilde{\mathbf{y}}^{\,v} = \sum_{u \in \mathcal{V}} K_h(v,u)\,\big(\mathbf{X}\boldsymbol{\beta}^u + \boldsymbol{\epsilon}^u\big) = \mathbf{X}\sum_{u \in \mathcal{V}} K_h(v,u)\,\boldsymbol{\beta}^u + \sum_{u \in \mathcal{V}} K_h(v,u)\,\boldsymbol{\epsilon}^u.$$

The GKS-based GLM estimator at $v$ is

$$\widehat{\boldsymbol{\beta}}^{\,v}_{\mathrm{GKS}} = (\mathbf{X}^\top\mathbf{X})^{-1}\mathbf{X}^\top\tilde{\mathbf{y}}^{\,v}.$$

Using the expression for $\tilde{\mathbf{y}}^{\,v}$ and the fact that $\mathbf{X}$ is fixed,

$$\widehat{\boldsymbol{\beta}}^{\,v}_{\mathrm{GKS}} = (\mathbf{X}^\top\mathbf{X})^{-1}\mathbf{X}^\top\left\{\mathbf{X}\sum_{u} K_h(v,u)\,\boldsymbol{\beta}^u + \sum_{u} K_h(v,u)\,\boldsymbol{\epsilon}^u\right\}.$$

Since $(\mathbf{X}^\top\mathbf{X})^{-1}\mathbf{X}^\top\mathbf{X} = \mathbf{I}_{KB}$, we obtain

$$\widehat{\boldsymbol{\beta}}^{\,v}_{\mathrm{GKS}} = \sum_{u \in \mathcal{V}} K_h(v,u)\,\boldsymbol{\beta}^u + (\mathbf{X}^\top\mathbf{X})^{-1}\mathbf{X}^\top\sum_{u \in \mathcal{V}} K_h(v,u)\,\boldsymbol{\epsilon}^u.$$

Taking expectation conditional on $\{\boldsymbol{\beta}^u\}_{u\in\mathcal{V}}$ and $\mathbf{X}$, and using $\mathbb{E}(\boldsymbol{\epsilon}^u) = \mathbf{0}$ for all $u$,

$$\mathbb{E}\Big(\widehat{\boldsymbol{\beta}}^{\,v}_{\mathrm{GKS}} \mid \{\boldsymbol{\beta}^u\}, \mathbf{X}\Big) = \sum_{u \in \mathcal{V}} K_h(v,u)\,\boldsymbol{\beta}^u,$$

which is Equation (1). Subtracting $\boldsymbol{\beta}^v$ from both sides yields the bias expression

$$\mathrm{Bias}\big(\widehat{\boldsymbol{\beta}}^{\,v}_{\mathrm{GKS}}\big) = \sum_{u \in \mathcal{V}} K_h(v,u)\,\boldsymbol{\beta}^u - \boldsymbol{\beta}^v.$$

In particular, the bias is zero if $\boldsymbol{\beta}^u$ is constant over all $u$ such that $K_h(v,u) > 0$, and is generally nonzero whenever $\boldsymbol{\beta}$ varies within the kernel support of $v$. □

**Proposition 2** (Variance attenuation under GKS-based GLM)**.** *In addition to the assumptions of Proposition 1, suppose that the temporal errors are independent across locations and share the same covariance* $\boldsymbol{\Sigma}_\epsilon$*.*

*Let*

$$\mathbf{R} = \mathbf{I}_T - \mathbf{X}(\mathbf{X}^\top\mathbf{X})^{-1}\mathbf{X}^\top$$

*be the GLM residual-forming matrix, and define*

$$\widehat{\boldsymbol{\epsilon}}^v_{\text{GKS}} = \mathbf{R}\,\tilde{\mathbf{y}}^v, \qquad \widehat{\boldsymbol{\epsilon}}^v = \mathbf{R}\,\mathbf{y}^v,$$

*as the GKS and unsmoothed GLM residuals, respectively.*

*Then the covariance of the GKS residuals satisfies*

$$\operatorname{Cov}(\widehat{\boldsymbol{\epsilon}}^v_{\text{GKS}}) = \left(\sum_{u\in\mathcal{V}} K_h(v,u)^2\right)\operatorname{Cov}(\widehat{\boldsymbol{\epsilon}}^v)\,, \tag{3}$$

*where* $\operatorname{Cov}(\widehat{\boldsymbol{\epsilon}}^v) = \mathbf{R}\boldsymbol{\Sigma}_\epsilon\mathbf{R}^\top$ *is the residual covariance in the absence of smoothing.*

*Moreover, if the kernel is nontrivial in the sense that there exist at least two locations* $u_1 \neq u_2$ *with* $K_h(v,u_j) > 0$ *for* $j = 1, 2$*, then*

$$0 < \sum_{u\in\mathcal{V}} K_h(v,u)^2 < 1,$$

*and hence*

$$\operatorname{Cov}(\widehat{\boldsymbol{\epsilon}}^v_{\text{GKS}}) \prec \operatorname{Cov}(\widehat{\boldsymbol{\epsilon}}^v)$$

*in strict Loewner order.*

*Proof.* Under the assumptions of Proposition 1 and the additional conditions of Proposition 2, write the noise at location $u$ as $\boldsymbol{\epsilon}^u$ with $\mathbb{E}(\boldsymbol{\epsilon}^u) = \mathbf{0}$ and $\operatorname{Cov}(\boldsymbol{\epsilon}^u) = \boldsymbol{\Sigma}_\epsilon$ for all $u \in \mathcal{V}$, and assume that $\boldsymbol{\epsilon}^u$ and $\boldsymbol{\epsilon}^{u'}$ are independent for $u \neq u'$.

For a fixed $v \in \mathcal{V}$, the smoothed noise contribution is

$$\tilde{\boldsymbol{\epsilon}}^v = \sum_{u\in\mathcal{V}} K_h(v,u)\,\boldsymbol{\epsilon}^u,$$

so $\mathbb{E}(\tilde{\boldsymbol{\epsilon}}^v) = \mathbf{0}$ and

$$\operatorname{Cov}(\tilde{\boldsymbol{\epsilon}}^v) = \mathbb{E}\big(\tilde{\boldsymbol{\epsilon}}^v\tilde{\boldsymbol{\epsilon}}^{v\top}\big) = \sum_{u,u'\in\mathcal{V}} K_h(v,u)K_h(v,u')\,\mathbb{E}\Big(\boldsymbol{\epsilon}^u\boldsymbol{\epsilon}^{u'\top}\Big)\,.$$

By spatial independence, $\mathbb{E}(\boldsymbol{\epsilon}^u \boldsymbol{\epsilon}^{u'\top}) = \mathbf{0}$ for $u \neq u'$, and $\mathbb{E}(\boldsymbol{\epsilon}^u \boldsymbol{\epsilon}^{u\top}) = \boldsymbol{\Sigma}_\epsilon$ for all $u$. Thus

$$\mathrm{Cov}(\tilde{\boldsymbol{\epsilon}}^v) = \sum_{u \in \mathcal{V}} K_h(v,u)^2 \, \boldsymbol{\Sigma}_\epsilon.$$

Let $\mathbf{R} = \mathbf{I}_T - \mathbf{X}(\mathbf{X}^\top \mathbf{X})^{-1} \mathbf{X}^\top$ denote the residual-forming matrix. For the unsmoothed GLM at location $v$,

$$\widehat{\boldsymbol{\epsilon}}^v = \mathbf{R}\,\mathbf{y}^v = \mathbf{R}\left(\mathbf{X}\boldsymbol{\beta}^v + \boldsymbol{\epsilon}^v\right) = \mathbf{R}\,\boldsymbol{\epsilon}^v$$

since $\mathbf{R}\mathbf{X} = \mathbf{0}$. Hence

$$\mathrm{Cov}(\widehat{\boldsymbol{\epsilon}}^v) = \mathbf{R}\,\boldsymbol{\Sigma}_\epsilon\,\mathbf{R}^\top.$$

For the GKS-based GLM, we similarly have

$$\widehat{\boldsymbol{\epsilon}}^v_{\mathrm{GKS}} = \mathbf{R}\,\tilde{\mathbf{y}}^v = \mathbf{R}\left(\mathbf{X}\tilde{\boldsymbol{\beta}}^v + \tilde{\boldsymbol{\epsilon}}^v\right) = \mathbf{R}\,\tilde{\boldsymbol{\epsilon}}^v,$$

where $\tilde{\boldsymbol{\beta}}^v = \sum_u K_h(v,u)\,\boldsymbol{\beta}^u$. Therefore

$$\mathrm{Cov}(\widehat{\boldsymbol{\epsilon}}^v_{\mathrm{GKS}}) = \mathbf{R}\,\mathrm{Cov}(\tilde{\boldsymbol{\epsilon}}^v)\,\mathbf{R}^\top = \left(\sum_{u \in \mathcal{V}} K_h(v,u)^2\right) \mathbf{R}\,\boldsymbol{\Sigma}_\epsilon\,\mathbf{R}^\top.$$

Using the expression for $\mathrm{Cov}(\widehat{\boldsymbol{\epsilon}}^v)$ above, this yields Equation (3):

$$\mathrm{Cov}(\widehat{\boldsymbol{\epsilon}}^v_{\mathrm{GKS}}) = \left(\sum_{u \in \mathcal{V}} K_h(v,u)^2\right) \mathrm{Cov}(\widehat{\boldsymbol{\epsilon}}^v)\,.$$

To establish the Loewner inequality, note that the weights satisfy $K_h(v,u) \geq 0$ and $\sum_u K_h(v,u) = 1$. Hence each weight lies in $[0,1]$, so $K_h(v,u)^2 \leq K_h(v,u)$ for every $u$, and therefore

$$\sum_{u \in \mathcal{V}} K_h(v,u)^2 \leq \sum_{u \in \mathcal{V}} K_h(v,u) = 1,$$

with equality if and only if all but at most one of the $K_h(v,u)$ are zero. Under the nontriviality assumption (at least two positive weights), we have $0 < \sum_u K_h(v,u)^2 < 1$.

Since $\mathrm{Cov}(\widehat{\boldsymbol{\epsilon}}^v)$ is positive semidefinite and $c \in (0,1)$ implies $cM \prec M$ for any positive definite matrix $M$, we conclude that

$$\mathrm{Cov}(\widehat{\boldsymbol{\epsilon}}^v_{\mathrm{GKS}}) \prec \mathrm{Cov}(\widehat{\boldsymbol{\epsilon}}^v)\,,$$

as claimed. □

**Corollary 1** (Variance underestimation under homoskedastic errors). *Under the setting of Proposition 2, suppose further that the temporal errors are homoskedastic and uncorrelated in time, with*

$$\mathbb{E}(\boldsymbol{\epsilon}^u) = \mathbf{0}, \qquad \mathrm{Cov}(\boldsymbol{\epsilon}^u) = \sigma^2 \mathbf{I}_T, \qquad u \in \mathcal{V}.$$

*Then*

$$\mathrm{Cov}(\widehat{\boldsymbol{\epsilon}}^v_{\mathrm{GKS}}) = \sigma^2 \left( \sum_{u \in \mathcal{V}} K_h(v,u)^2 \right) \mathbf{R}\mathbf{R}^\top.$$

*In particular, if $\hat{\sigma}^2$ denotes the usual unbiased estimator from the unsmoothed GLM,*

$$\hat{\sigma}^2 = \frac{\|\widehat{\boldsymbol{\epsilon}}^v\|^2}{T - KB},$$

*then the analogous estimator based on GKS residuals,*

$$\hat{\sigma}^2_{\mathrm{GKS}} = \frac{\|\widehat{\boldsymbol{\epsilon}}^v_{\mathrm{GKS}}\|^2}{T - KB},$$

*satisfies*

$$\mathbb{E}\big(\hat{\sigma}^2_{\mathrm{GKS}}\big) = \sigma^2 \left( \sum_{u \in \mathcal{V}} K_h(v,u)^2 \right) \; < \; \sigma^2.$$

*Thus, GKS always induces downward bias in the usual variance estimator, by an attenuation factor between* 0 *and* 1 *determined by the kernel weights.*

*Proof.* Under the additional assumption that the temporal errors are homoskedastic and uncorrelated in time, we have

$$\mathbb{E}(\boldsymbol{\epsilon}^u) = \mathbf{0}, \qquad \mathrm{Cov}(\boldsymbol{\epsilon}^u) = \sigma^2 \mathbf{I}_T \quad \text{for all } u \in \mathcal{V},$$

so in Proposition 2 the common covariance matrix is $\boldsymbol{\Sigma}_\epsilon = \sigma^2 \mathbf{I}_T$. Thus the unsmoothed GLM residual covariance becomes

$$\mathrm{Cov}(\widehat{\boldsymbol{\epsilon}}^v) = \mathbf{R}\boldsymbol{\Sigma}_\epsilon \mathbf{R}^\top = \sigma^2 \, \mathbf{R}\mathbf{R}^\top.$$

By Proposition 2,

$$\mathrm{Cov}(\widehat{\boldsymbol{\epsilon}}^v_{\mathrm{GKS}}) = \left( \sum_{u \in \mathcal{V}} K_h(v,u)^2 \right) \mathrm{Cov}(\widehat{\boldsymbol{\epsilon}}^v)\,,$$

and substituting the expression above yields

$$\mathrm{Cov}(\widehat{\boldsymbol{\epsilon}}^v_{\mathrm{GKS}}) = \sigma^2 \left( \sum_{u \in \mathcal{V}} K_h(v,u)^2 \right) \mathbf{R}\mathbf{R}^\top.$$

In our GLM with $KB$ regressors (assumed full rank), the usual unbiased estimator of $\sigma^2$ from the unsmoothed GLM is

$$\hat{\sigma}^2 = \frac{\|\widehat{\boldsymbol{\epsilon}}^v\|^2}{T - KB},$$

since the residual-maker matrix $\mathbf{R}$ is symmetric idempotent with $\mathrm{tr}(\mathbf{R}\mathbf{R}^\top) = \mathrm{tr}(\mathbf{R}) = T - KB$. Using

$$\mathbb{E}\big(\|\widehat{\boldsymbol{\epsilon}}^v\|^2\big) = \mathrm{tr}(\mathrm{Cov}(\widehat{\boldsymbol{\epsilon}}^v)) = \mathrm{tr}\big(\sigma^2\, \mathbf{R}\mathbf{R}^\top\big) = \sigma^2(T - KB),$$

we obtain $\mathbb{E}(\hat{\sigma}^2) = \sigma^2$.

For the GKS-based estimator,

$$\hat{\sigma}^2_{\mathrm{GKS}} = \frac{\|\widehat{\boldsymbol{\epsilon}}^v_{\mathrm{GKS}}\|^2}{T - KB},$$

and

$$\mathbb{E}\big(\|\widehat{\boldsymbol{\epsilon}}^v_{\mathrm{GKS}}\|^2\big) = \mathrm{tr}(\mathrm{Cov}(\widehat{\boldsymbol{\epsilon}}^v_{\mathrm{GKS}})) = \sigma^2 \left( \sum_{u \in \mathcal{V}} K_h(v,u)^2 \right) \mathrm{tr}(\mathbf{R}\mathbf{R}^\top) = \sigma^2 \left( \sum_{u \in \mathcal{V}} K_h(v,u)^2 \right) (T - KB).$$

Dividing by $T - KB$ gives

$$\mathbb{E}\big(\hat{\sigma}^2_{\mathrm{GKS}}\big) = \sigma^2 \left( \sum_{u \in \mathcal{V}} K_h(v,u)^2 \right).$$

Since the kernel weights are nonnegative, sum to 1, and are nontrivial (i.e., there exists at least one $u \neq v$ with $K_h(v,u) > 0$), we have $0 < \sum_{u \in \mathcal{V}} K_h(v,u)^2 < 1$. Therefore,

$$\mathbb{E}\big(\hat{\sigma}^2_{\mathrm{GKS}}\big) < \sigma^2,$$

so GKS always induces downward bias in the usual variance estimator, by an attenuation factor between 0 and 1 determined by the kernel weights. □

**Theorem 1** (Consistency of SPLASH activation estimates). *Fix a parcel and consider the linear model*

$$\mathbf{Y} = \mathbf{Z}\mathbf{P}\boldsymbol{\Phi}\,\boldsymbol{\Gamma} + \boldsymbol{\epsilon}, \qquad \mathbb{E}(\boldsymbol{\epsilon}) = \mathbf{0},\ \mathrm{Cov}(\boldsymbol{\epsilon}) = \boldsymbol{\Sigma},$$

*where* $\mathbf{Y} \in \mathbb{R}^{TV}$ *stacks all observations in the parcel, the design matrix*

$$\mathbf{A} = \mathbf{Z}\mathbf{P}\boldsymbol{\Phi} \in \mathbb{R}^{(TV)\times(KBM)}$$

*has full column rank, and* $\boldsymbol{\Gamma} \in \mathbb{R}^{KBM}$ *is the vector of spatial-spline coefficients.*

*Because the block-diagonal noise covariance* $\boldsymbol{\Sigma}$ *is unknown, SPLASH uses a feasible generalized least squares (FGLS) estimator in which* $\boldsymbol{\Sigma}$ *is replaced by a consistent estimator* $\hat{\boldsymbol{\Sigma}}$*. The estimator is*

$$\hat{\boldsymbol{\Gamma}} = \left(\mathbf{A}^\top \hat{\boldsymbol{\Sigma}}^{-1} \mathbf{A}\right)^{-1} \mathbf{A}^\top \hat{\boldsymbol{\Sigma}}^{-1} \mathbf{Y}.$$

*Assume the following:*

1. *The number of independent experimental units* $N$ *(e.g., subjects) tends to infinity, while the dimensions* $TV$ *and* $KBM$ *remain fixed.*
2. *The noise covariance estimator is consistent:*

$$\hat{\boldsymbol{\Sigma}}^{-1} \xrightarrow{p} \boldsymbol{\Sigma}^{-1} \qquad \text{as } N \to \infty.$$

3. $\mathbf{A}$ *has full column rank, so that* $\mathbf{A}^\top \boldsymbol{\Sigma}^{-1} \mathbf{A}$ *is positive definite.*

*Then the feasible GLS estimator is consistent:*

$$\hat{\boldsymbol{\Gamma}} \xrightarrow{p} \boldsymbol{\Gamma} \qquad \text{as } N \to \infty.$$

*Finally, SPLASH reconstructs location-specific activation coefficients using*

$$\hat{\boldsymbol{\beta}}_{\mathrm{sp}} = \mathbf{P}\left(\mathbf{I}_{KB} \otimes \mathbf{S}\right)\hat{\boldsymbol{\Gamma}},$$

*and by continuity of this mapping,*

$$\hat{\boldsymbol{\beta}}_{\mathrm{sp}} \xrightarrow{p} \boldsymbol{\beta}_{\mathrm{sp}} \qquad \text{as } N \to \infty.$$

*Proof.* For clarity, we sketch the argument in the balanced case where each of the $N$ independent experimental units (e.g., participants) contributes data following the same parcel-wise SPLASH model

$$\mathbf{Y}_i = \mathbf{A}\,\boldsymbol{\Gamma} + \boldsymbol{\epsilon}_i, \qquad i = 1, \ldots, N,$$

with $\mathbb{E}(\boldsymbol{\epsilon}_i) = \mathbf{0}$ and $\mathrm{Cov}(\boldsymbol{\epsilon}_i) = \boldsymbol{\Sigma}$, and the $\boldsymbol{\epsilon}_i$ are independent across $i$. Here $\mathbf{Y}_i \in \mathbb{R}^{TV}$ and $\mathbf{A} \in \mathbb{R}^{(TV)\times(KBM)}$ has full column rank.

Stacking all $N$ observations, define

$$\mathbf{Y}_N = \begin{bmatrix} \mathbf{Y}_1 \\ \vdots \\ \mathbf{Y}_N \end{bmatrix} \in \mathbb{R}^{(TV)N}, \qquad \mathbf{A}_N = \mathbf{1}_N \otimes \mathbf{A} \in \mathbb{R}^{(TV)N\times(KBM)}, \qquad \boldsymbol{\epsilon}_N = \begin{bmatrix} \boldsymbol{\epsilon}_1 \\ \vdots \\ \boldsymbol{\epsilon}_N \end{bmatrix},$$

so that

$$\mathbf{Y}_N = \mathbf{A}_N\boldsymbol{\Gamma} + \boldsymbol{\epsilon}_N, \quad \mathrm{Cov}(\boldsymbol{\epsilon}_N) = \boldsymbol{\Sigma}_N = \mathbf{I}_N \otimes \boldsymbol{\Sigma}.$$

If $\boldsymbol{\Sigma}_N$ were known, the GLS estimator would be

$$\hat{\boldsymbol{\Gamma}}_{\mathrm{GLS},N} = \left(\mathbf{A}_N^\top \boldsymbol{\Sigma}_N^{-1} \mathbf{A}_N\right)^{-1} \mathbf{A}_N^\top \boldsymbol{\Sigma}_N^{-1} \mathbf{Y}_N.$$

Using properties of the Kronecker product,

$$\mathbf{A}_N^\top \boldsymbol{\Sigma}_N^{-1} \mathbf{A}_N = (\mathbf{1}_N^\top \mathbf{1}_N) \otimes (\mathbf{A}^\top \boldsymbol{\Sigma}^{-1} \mathbf{A}) = N\, \mathbf{A}^\top \boldsymbol{\Sigma}^{-1} \mathbf{A},$$

and

$$\mathbf{A}_N^\top \boldsymbol{\Sigma}_N^{-1} \boldsymbol{\epsilon}_N = \sum_{i=1}^{N} \mathbf{A}^\top \boldsymbol{\Sigma}^{-1} \boldsymbol{\epsilon}_i.$$

Hence

$$\begin{aligned}
\hat{\boldsymbol{\Gamma}}_{\mathrm{GLS},N} &= \left(N\, \mathbf{A}^\top \boldsymbol{\Sigma}^{-1} \mathbf{A}\right)^{-1} \mathbf{A}_N^\top \boldsymbol{\Sigma}_N^{-1} (\mathbf{A}_N \boldsymbol{\Gamma} + \boldsymbol{\epsilon}_N) \\
&= \boldsymbol{\Gamma} + \left(N\, \mathbf{A}^\top \boldsymbol{\Sigma}^{-1} \mathbf{A}\right)^{-1} \mathbf{A}_N^\top \boldsymbol{\Sigma}_N^{-1} \boldsymbol{\epsilon}_N \\
&= \boldsymbol{\Gamma} + \left(\mathbf{A}^\top \boldsymbol{\Sigma}^{-1} \mathbf{A}\right)^{-1} \frac{1}{N} \sum_{i=1}^{N} \mathbf{A}^\top \boldsymbol{\Sigma}^{-1} \boldsymbol{\epsilon}_i.
\end{aligned}$$

The summands $\mathbf{A}^\top \boldsymbol{\Sigma}^{-1} \boldsymbol{\epsilon}_i$ are i.i.d. with mean $\mathbf{0}$ and covariance

$$\mathbf{A}^\top \boldsymbol{\Sigma}^{-1} \boldsymbol{\Sigma} \boldsymbol{\Sigma}^{-1} \mathbf{A} = \mathbf{A}^\top \boldsymbol{\Sigma}^{-1} \mathbf{A}.$$

By the multivariate law of large numbers,

$$\frac{1}{N} \sum_{i=1}^{N} \mathbf{A}^\top \boldsymbol{\Sigma}^{-1} \boldsymbol{\epsilon}_i \xrightarrow{p} \mathbf{0} \quad \text{as } N \to \infty.$$

Since $\mathbf{A}^\top \boldsymbol{\Sigma}^{-1} \mathbf{A}$ is positive definite by assumption,

$$\hat{\boldsymbol{\Gamma}}_{\mathrm{GLS},N} \xrightarrow{p} \boldsymbol{\Gamma}.$$

Now consider the feasible GLS estimator

$$\hat{\boldsymbol{\Gamma}}_N = \left(\mathbf{A}_N^\top \hat{\boldsymbol{\Sigma}}_N^{-1} \mathbf{A}_N\right)^{-1} \mathbf{A}_N^\top \hat{\boldsymbol{\Sigma}}_N^{-1} \mathbf{Y}_N,$$

where $\hat{\boldsymbol{\Sigma}}_N^{-1}$ is a consistent estimator of $\boldsymbol{\Sigma}_N^{-1}$ in the sense that

$$\hat{\boldsymbol{\Sigma}}_N^{-1} \xrightarrow{p} \boldsymbol{\Sigma}_N^{-1} \qquad \text{as } N \to \infty.$$

The map

$$(\mathbf{M}, \mathbf{y}) \mapsto \left(\mathbf{A}_N^\top \mathbf{M} \mathbf{A}_N\right)^{-1} \mathbf{A}_N^\top \mathbf{M} \mathbf{y}$$

is continuous on the set where $\mathbf{A}_N^\top \mathbf{M} \mathbf{A}_N$ is invertible, which occurs with probability tending to 1 as $\hat{\boldsymbol{\Sigma}}_N^{-1}$ approaches $\boldsymbol{\Sigma}_N^{-1}$. By the continuous mapping theorem and Slutsky's theorem,

$$\hat{\boldsymbol{\Gamma}}_N - \hat{\boldsymbol{\Gamma}}_{\mathrm{GLS},N} \xrightarrow{p} \mathbf{0}.$$

Combining with $\hat{\boldsymbol{\Gamma}}_{\mathrm{GLS},N} \xrightarrow{p} \boldsymbol{\Gamma}$ gives

$$\hat{\boldsymbol{\Gamma}}_N \xrightarrow{p} \boldsymbol{\Gamma}.$$

Finally, SPLASH computes activation estimates via the fixed linear map

$$\hat{\boldsymbol{\beta}}_{\mathrm{sp},N} = \mathbf{P}(\mathbf{I}_{KB} \otimes \mathbf{S})\, \hat{\boldsymbol{\Gamma}}_N.$$

Since this transformation is continuous, another application of the continuous mapping theorem yields

$$\hat{\boldsymbol{\beta}}_{\mathrm{sp},N} \xrightarrow{p} \mathbf{P}(\mathbf{I}_{KB} \otimes \mathbf{S})\, \boldsymbol{\Gamma} = \boldsymbol{\beta}_{\mathrm{sp}},$$

as claimed. □

**Corollary 2** (Unbiasedness under independent homoskedastic errors)**.** *In the setting of Theorem 1, suppose in addition that the noise is independent and homoskedastic, so that*

$$\mathbb{E}(\boldsymbol{\epsilon}) = \mathbf{0}, \qquad \mathrm{Cov}(\boldsymbol{\epsilon}) = \boldsymbol{\Sigma} = \sigma^2 \mathbf{I}_{TV} \quad \textit{for some } \sigma^2 > 0,$$

*and that the working covariance used in FGLS is restricted to the same spherical form*

$$\hat{\boldsymbol{\Sigma}} = \hat{\sigma}^2 \mathbf{I}_{TV}.$$

*Then the FGLS estimator reduces to ordinary least squares,*

$$\hat{\boldsymbol{\Gamma}} = \left(\mathbf{A}^\top \mathbf{A}\right)^{-1} \mathbf{A}^\top \mathbf{Y},$$

*and is unbiased:*

$$\mathbb{E}(\hat{\boldsymbol{\Gamma}}) = \boldsymbol{\Gamma}.$$

*Moreover, with the SPLASH reconstruction*

$$\hat{\boldsymbol{\beta}}_{\text{sp}} = \mathbf{P}(\mathbf{I}_{KB} \otimes \mathbf{S})\, \hat{\boldsymbol{\Gamma}},$$

*we have*

$$\mathbb{E}(\hat{\boldsymbol{\beta}}_{\text{sp}}) = \boldsymbol{\beta}_{\text{sp}}.$$

*Proof.* Under the additional assumptions, $\boldsymbol{\Sigma} = \sigma^2 \mathbf{I}_{TV}$ and $\hat{\boldsymbol{\Sigma}} = \hat{\sigma}^2 \mathbf{I}_{TV}$. Substituting this working covariance into the FGLS estimator in Theorem 1 gives

$$\hat{\boldsymbol{\Gamma}} = \left(\mathbf{A}^\top \hat{\boldsymbol{\Sigma}}^{-1} \mathbf{A}\right)^{-1} \mathbf{A}^\top \hat{\boldsymbol{\Sigma}}^{-1} \mathbf{Y} = \left(\hat{\sigma}^{-2} \mathbf{A}^\top \mathbf{A}\right)^{-1} \hat{\sigma}^{-2} \mathbf{A}^\top \mathbf{Y} = \left(\mathbf{A}^\top \mathbf{A}\right)^{-1} \mathbf{A}^\top \mathbf{Y},$$

so FGLS coincides exactly with ordinary least squares in the linear model $\mathbf{Y} = \mathbf{A}\boldsymbol{\Gamma} + \boldsymbol{\epsilon}$ with $\mathbb{E}(\boldsymbol{\epsilon}) = \mathbf{0}$. By standard OLS theory,

$$\mathbb{E}(\hat{\boldsymbol{\Gamma}}) = \left(\mathbf{A}^\top \mathbf{A}\right)^{-1} \mathbf{A}^\top \mathbb{E}(\mathbf{Y}) = \left(\mathbf{A}^\top \mathbf{A}\right)^{-1} \mathbf{A}^\top \mathbf{A} \boldsymbol{\Gamma} = \boldsymbol{\Gamma}.$$

The SPLASH activation estimates are obtained via the fixed linear map $\hat{\boldsymbol{\beta}}_{\text{sp}} = \mathbf{P}(\mathbf{I}_{KB} \otimes \mathbf{S})\, \hat{\boldsymbol{\Gamma}}$, so taking expectations and using linearity yields

$$\mathbb{E}(\hat{\boldsymbol{\beta}}_{\text{sp}}) = \mathbf{P}(\mathbf{I}_{KB} \otimes \mathbf{S})\, \mathbb{E}(\hat{\boldsymbol{\Gamma}}) = \mathbf{P}(\mathbf{I}_{KB} \otimes \mathbf{S})\, \boldsymbol{\Gamma} = \boldsymbol{\beta}_{\text{sp}}.$$

□

**Theorem 2** (Consistency of SPLASH variance estimation)**.** *Under the assumptions of Theorem 1, let*

$$\widehat{\mathrm{Var}}(\hat{\boldsymbol{\Gamma}}) = \left(\mathbf{A}^\top \hat{\boldsymbol{\Sigma}}^{-1} \mathbf{A}\right)^{-1}$$

*denote the usual plug-in covariance estimator for the FGLS estimator* $\hat{\boldsymbol{\Gamma}}$*.*

*Then, as* $N \to \infty$*,*

$$\widehat{\mathrm{Var}}(\hat{\boldsymbol{\Gamma}}) \xrightarrow{p} \left(\mathbf{A}^\top \boldsymbol{\Sigma}^{-1} \mathbf{A}\right)^{-1} = \mathrm{Var}(\hat{\boldsymbol{\Gamma}}).$$

*Moreover, for any fixed linear transformation* $\mathbf{L} \in \mathbb{R}^{r \times KBM}$*, the covariance estimator*

$$\widehat{\mathrm{Var}}(\mathbf{L}\hat{\boldsymbol{\Gamma}}) = \mathbf{L}\, \widehat{\mathrm{Var}}(\hat{\boldsymbol{\Gamma}})\, \mathbf{L}^\top$$

*satisfies*

$$\widehat{\mathrm{Var}}(\mathbf{L}\hat{\boldsymbol{\Gamma}}) \xrightarrow{p} \mathrm{Var}(\mathbf{L}\hat{\boldsymbol{\Gamma}}).$$

*In particular, taking* $\mathbf{L} = \mathbf{P}(\mathbf{I}_{KB} \otimes \mathbf{S})$ *yields*

$$\widehat{\mathrm{Var}}(\hat{\boldsymbol{\beta}}_{\mathrm{sp}}) \xrightarrow{p} \mathrm{Var}(\hat{\boldsymbol{\beta}}_{\mathrm{sp}}),$$

*so SPLASH provides asymptotically valid uncertainty quantification for the smoothed activation field.*

*Proof.* Under the stacked model in the proof of Theorem 1, the GLS covariance of $\hat{\boldsymbol{\Gamma}}_{\mathrm{GLS},N}$ is

$$\mathrm{Var}(\hat{\boldsymbol{\Gamma}}_{\mathrm{GLS},N}) = \left(\mathbf{A}_N^\top \boldsymbol{\Sigma}_N^{-1} \mathbf{A}_N\right)^{-1} = \left(N\, \mathbf{A}^\top \boldsymbol{\Sigma}^{-1} \mathbf{A}\right)^{-1} = \frac{1}{N} \left(\mathbf{A}^\top \boldsymbol{\Sigma}^{-1} \mathbf{A}\right)^{-1}.$$

The feasible covariance estimator is

$$\widehat{\mathrm{Var}}(\hat{\boldsymbol{\Gamma}}_N) = \left(\mathbf{A}_N^\top \hat{\boldsymbol{\Sigma}}_N^{-1} \mathbf{A}_N\right)^{-1}.$$

By assumption, $\hat{\boldsymbol{\Sigma}}_N^{-1} \xrightarrow{p} \boldsymbol{\Sigma}_N^{-1}$, and the map

$$\mathbf{M} \mapsto \left(\mathbf{A}_N^\top \mathbf{M} \mathbf{A}_N\right)^{-1}$$

is continuous on the set of matrices for which $\mathbf{A}_N^\top \mathbf{M} \mathbf{A}_N$ is invertible, which holds with probability tending to 1. By the continuous mapping theorem,

$$\widehat{\mathrm{Var}}(\hat{\boldsymbol{\Gamma}}_N) \xrightarrow{p} \left(\mathbf{A}_N^\top \boldsymbol{\Sigma}_N^{-1} \mathbf{A}_N\right)^{-1} = \mathrm{Var}(\hat{\boldsymbol{\Gamma}}_{\mathrm{GLS},N}).$$

The feasible GLS estimator $\hat{\boldsymbol{\Gamma}}_N$ is asymptotically equivalent to the GLS estimator $\hat{\boldsymbol{\Gamma}}_{\mathrm{GLS},N}$, in the sense that $\hat{\boldsymbol{\Gamma}}_N - \hat{\boldsymbol{\Gamma}}_{\text{GLS},N} \xrightarrow{p} \mathbf{0}$. Combining this with the convergence of the covariance estimator and the fact that $\text{Var}(\hat{\boldsymbol{\Gamma}}_N) \to \text{Var}(\hat{\boldsymbol{\Gamma}}_{\text{GLS},N})$ as $N \to \infty$, we obtain

$$\widehat{\text{Var}}(\hat{\boldsymbol{\Gamma}}_N) \xrightarrow{p} \text{Var}(\hat{\boldsymbol{\Gamma}}_N).$$

For any fixed linear transformation $\mathbf{L}$, we have

$$\widehat{\text{Var}}(\mathbf{L}\hat{\boldsymbol{\Gamma}}_N) = \mathbf{L}\,\widehat{\text{Var}}(\hat{\boldsymbol{\Gamma}}_N)\,\mathbf{L}^\top \xrightarrow{p} \mathbf{L}\,\text{Var}(\hat{\boldsymbol{\Gamma}}_N)\,\mathbf{L}^\top = \text{Var}(\mathbf{L}\hat{\boldsymbol{\Gamma}}_N),$$

again by continuity of the map $M \mapsto LML^\top$. Taking $\mathbf{L} = \mathbf{P}(\mathbf{I}_{KB} \otimes \mathbf{S})$ yields

$$\widehat{\text{Var}}(\hat{\boldsymbol{\beta}}_{\text{sp},N}) \xrightarrow{p} \text{Var}(\hat{\boldsymbol{\beta}}_{\text{sp},N}),$$

which is the desired result. □

**Corollary 3** (Unbiased variance under independent errors)**.** *In the setting of Corollary 2, suppose the error variance $\sigma^2$ is estimated by*

$$\hat{\sigma}^2 = \frac{\|\mathbf{Y} - \mathbf{A}\hat{\boldsymbol{\Gamma}}\|^2}{TV - KBM},$$

*and define*

$$\widehat{\text{Var}}(\hat{\boldsymbol{\Gamma}}) = \hat{\sigma}^2\,(\mathbf{A}^\top\mathbf{A})^{-1}, \qquad \widehat{\text{Var}}(\hat{\boldsymbol{\beta}}_{\text{sp}}) = \mathbf{L}\,\widehat{\text{Var}}(\hat{\boldsymbol{\Gamma}})\,\mathbf{L}^\top,$$

*with $\mathbf{L} = \mathbf{P}(\mathbf{I}_{KB} \otimes \mathbf{S})$ as above.*

*Then*

$$\mathbb{E}\Big(\widehat{\text{Var}}(\hat{\boldsymbol{\Gamma}})\Big) = \text{Var}(\hat{\boldsymbol{\Gamma}}), \qquad \mathbb{E}\Big(\widehat{\text{Var}}(\hat{\boldsymbol{\beta}}_{\text{sp}})\Big) = \text{Var}(\hat{\boldsymbol{\beta}}_{\text{sp}}),$$

*so SPLASH achieves finite-sample unbiased variance estimation under independent errors.*

*Proof.* In the independent homoskedastic setting of Corollary 2, FGLS reduces to ordinary least squares, and the usual unbiased estimator of $\sigma^2$ is

$$\hat{\sigma}^2 = \frac{\|\mathbf{Y} - \mathbf{A}\hat{\boldsymbol{\Gamma}}\|^2}{TV - KBM}, \qquad \mathbb{E}(\hat{\sigma}^2) = \sigma^2.$$

The corresponding covariance estimator for the spline coefficients is

$$\widehat{\text{Var}}(\hat{\boldsymbol{\Gamma}}) = \hat{\sigma}^2\,(\mathbf{A}^\top\mathbf{A})^{-1}.$$

Taking expectations and using $\mathbb{E}(\hat{\sigma}^2) = \sigma^2$ gives

$$\mathbb{E}\Big(\widehat{\mathrm{Var}}(\hat{\boldsymbol{\Gamma}})\Big) = \sigma^2 \, (\mathbf{A}^\top \mathbf{A})^{-1} = \mathrm{Var}(\hat{\boldsymbol{\Gamma}}),$$

so the estimator is exactly unbiased for the true covariance of $\hat{\boldsymbol{\Gamma}}$.

For any fixed linear transformation $\mathbf{L}$,

$$\widehat{\mathrm{Var}}(\mathbf{L}\hat{\boldsymbol{\Gamma}}) = \mathbf{L}\, \widehat{\mathrm{Var}}(\hat{\boldsymbol{\Gamma}})\, \mathbf{L}^\top.$$

Taking expectations yields

$$\mathbb{E}\Big(\widehat{\mathrm{Var}}(\mathbf{L}\hat{\boldsymbol{\Gamma}})\Big) = \mathbf{L}\, \mathrm{Var}(\hat{\boldsymbol{\Gamma}})\, \mathbf{L}^\top = \mathrm{Var}(\mathbf{L}\hat{\boldsymbol{\Gamma}}).$$

With $\mathbf{L} = \mathbf{P}(\mathbf{I}_{KB} \otimes \mathbf{S})$, this gives

$$\mathbb{E}\Big(\widehat{\mathrm{Var}}(\hat{\boldsymbol{\beta}}_{\mathrm{sp}})\Big) = \mathrm{Var}(\hat{\boldsymbol{\beta}}_{\mathrm{sp}}).$$

Thus SPLASH achieves finite-sample unbiased variance estimation for both the spline coefficients and the reconstructed activation field under independent errors. □

### 2 Additional Simulation Results

This section presents supplementary simulation results complementing those in Section 3.2. Our full study evaluates SPLASH and competing methods across three SNR regimes (High, Moderate, and Low) and two task paradigms (Task 1 and Task 2). The main paper reports the primary High-SNR results for Task 1. Here, we provide the additional estimation and inference results for the Moderate- and Low-SNR settings, as well as the corresponding results for Task 2 under High SNR. For completeness, we also include the mean squared error (MSE) for each method across all scenarios.

Across all three SNR settings, SPLASH consistently produces clear, non-oversmoothed activation coefficient maps and well-localized activation regions via selective inference. SPLASH also achieves the strongest overall performance in terms of MSE, FPR, and FNR.

#### 2.1 High-SNR Setting

##### 2.1.1 Task 2

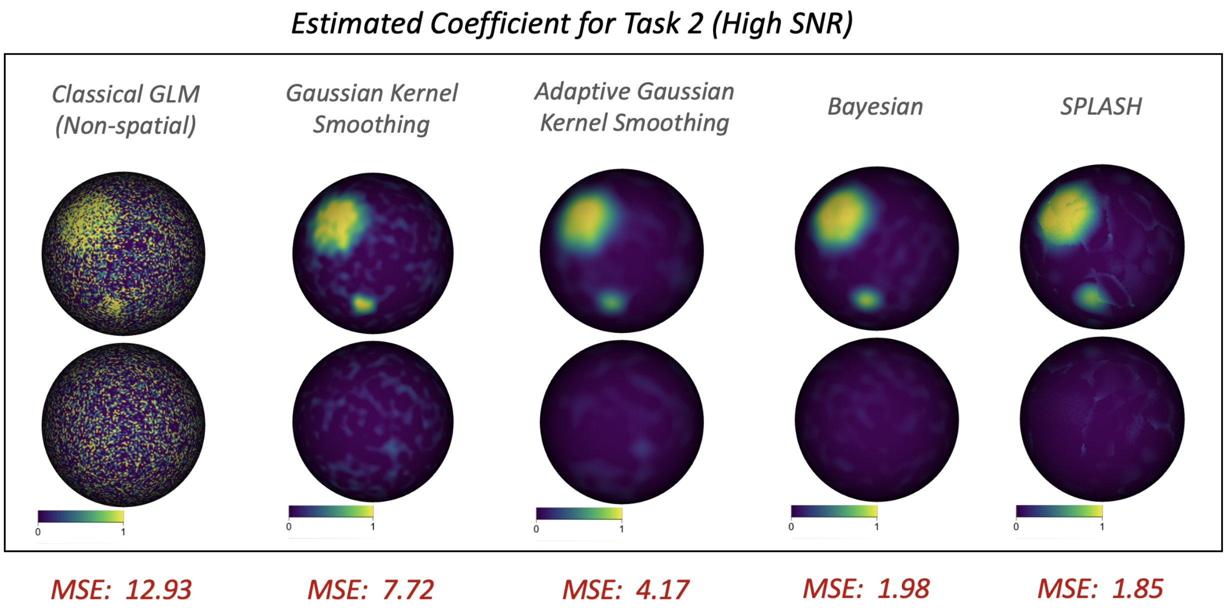

Figure S1: Estimated Task 2 coefficients under the High-SNR setting. MSE values are shown in red.

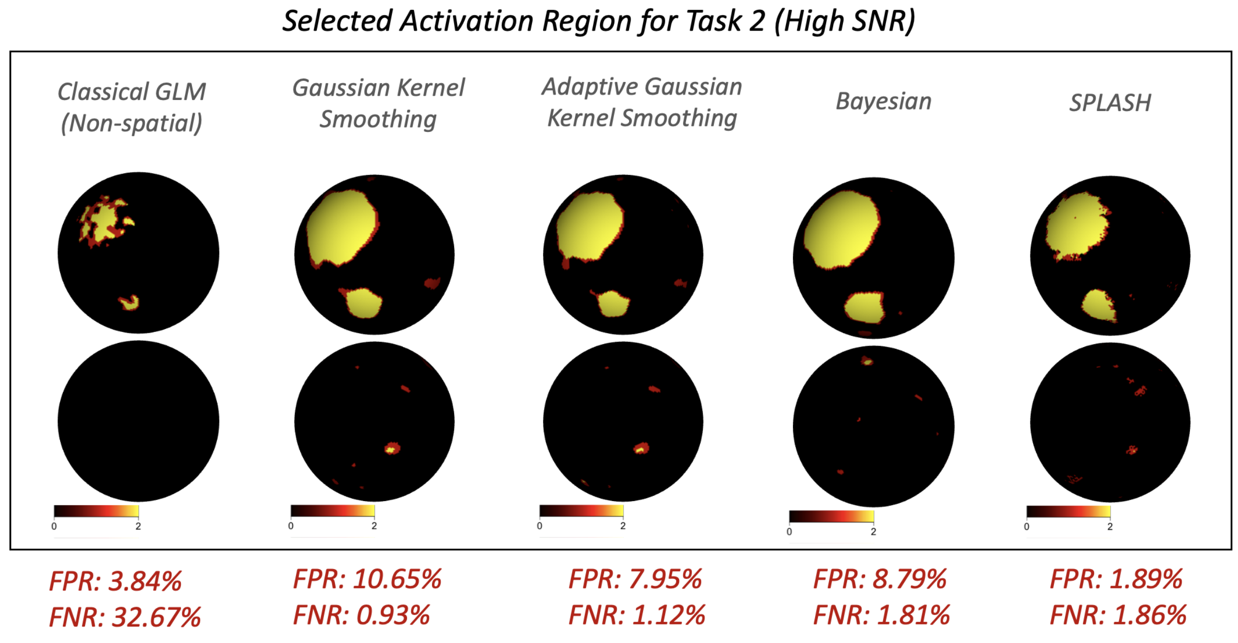

Figure S2: Detected activation for Task 2 under the high-SNR setting. False positive (FPR) and false negative (FNR) rates are shown in red.

#### 2.2 Moderate-SNR Setting

##### 2.2.1 Task 1

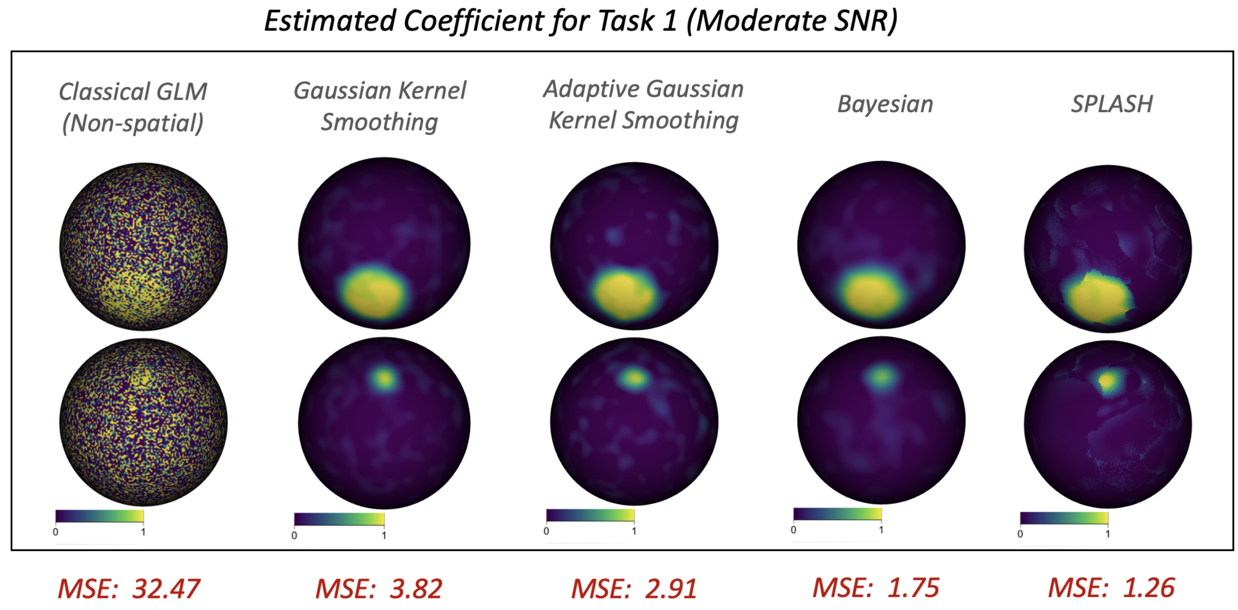

Figure S3: Estimated Task 1 coefficients under the moderate-SNR setting. MSE values are shown in red.

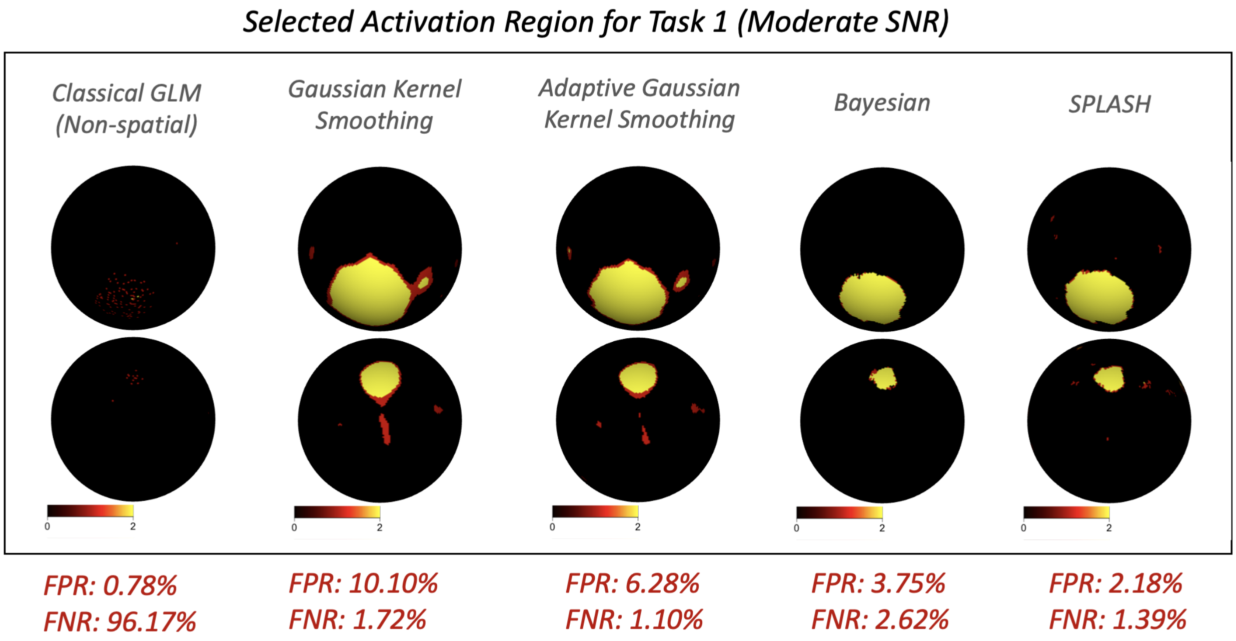

Figure S4: Detected activation for Task 1 under the moderate-SNR setting. False positive (FPR) and false negative (FNR) rates are shown in red.

##### 2.2.2 Task 2

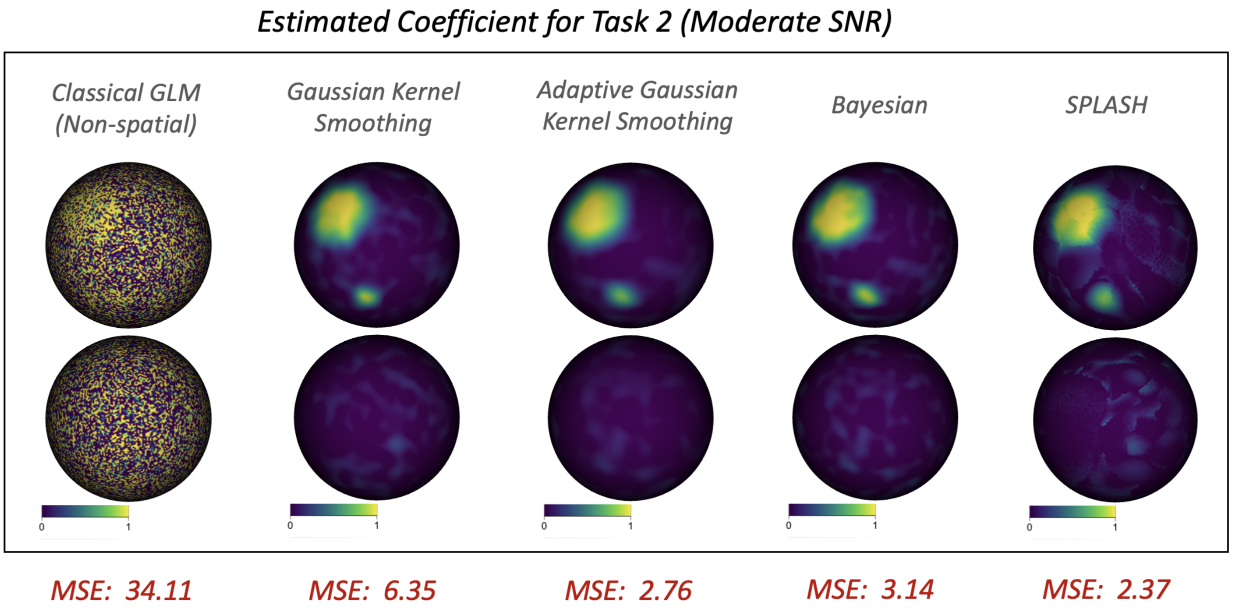

Figure S5: Estimated Task 2 coefficients under the moderate-SNR setting. MSE values are shown in red.

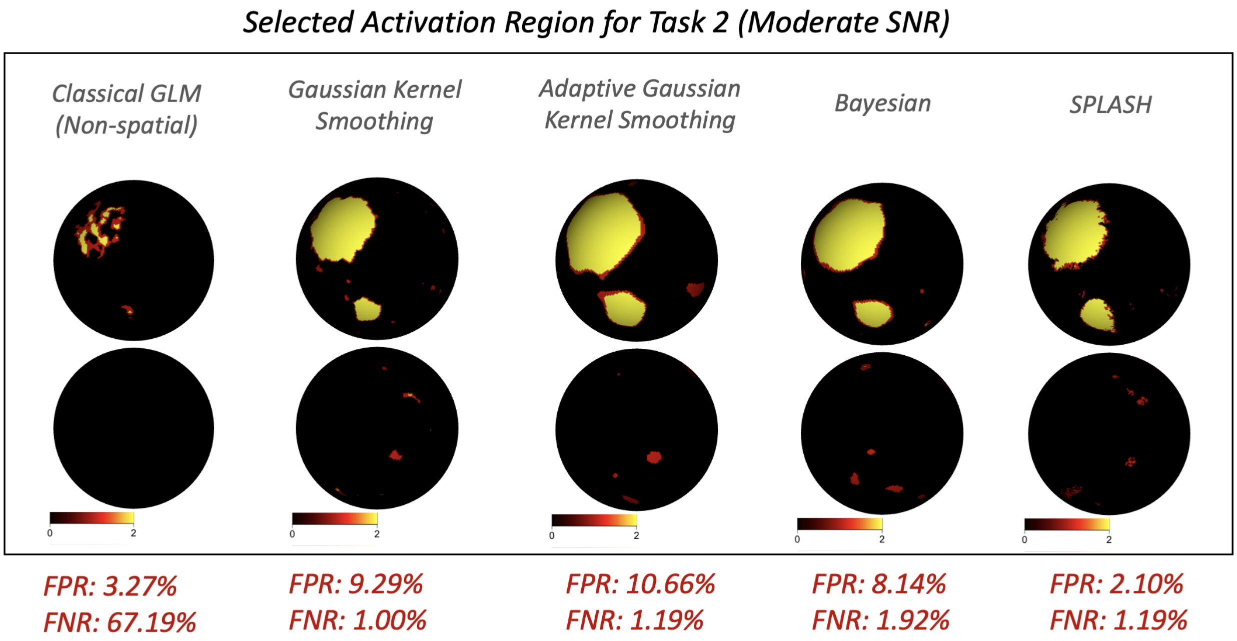

Figure S6: Detected activation for Task 2 under the moderate-SNR setting. False positive (FPR) and false negative (FNR) rates are shown in red.

#### 2.3 Low-SNR Setting

##### 2.3.1 Task 1

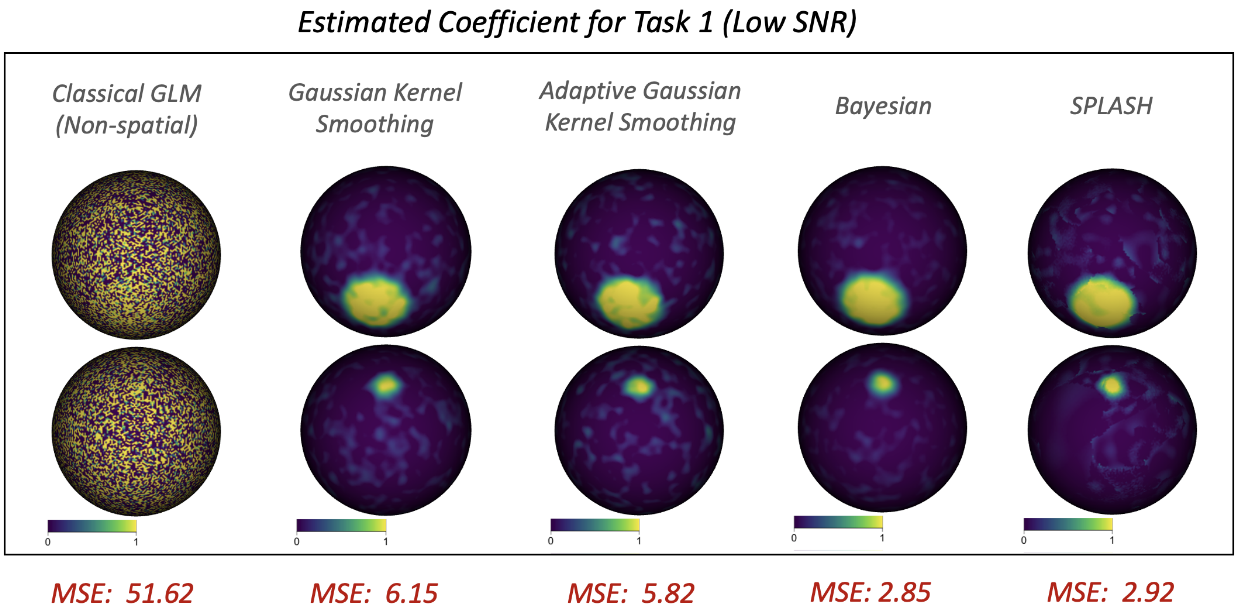

Figure S7: Estimated Task 1 coefficients under the low-SNR setting. MSE values are shown in red.

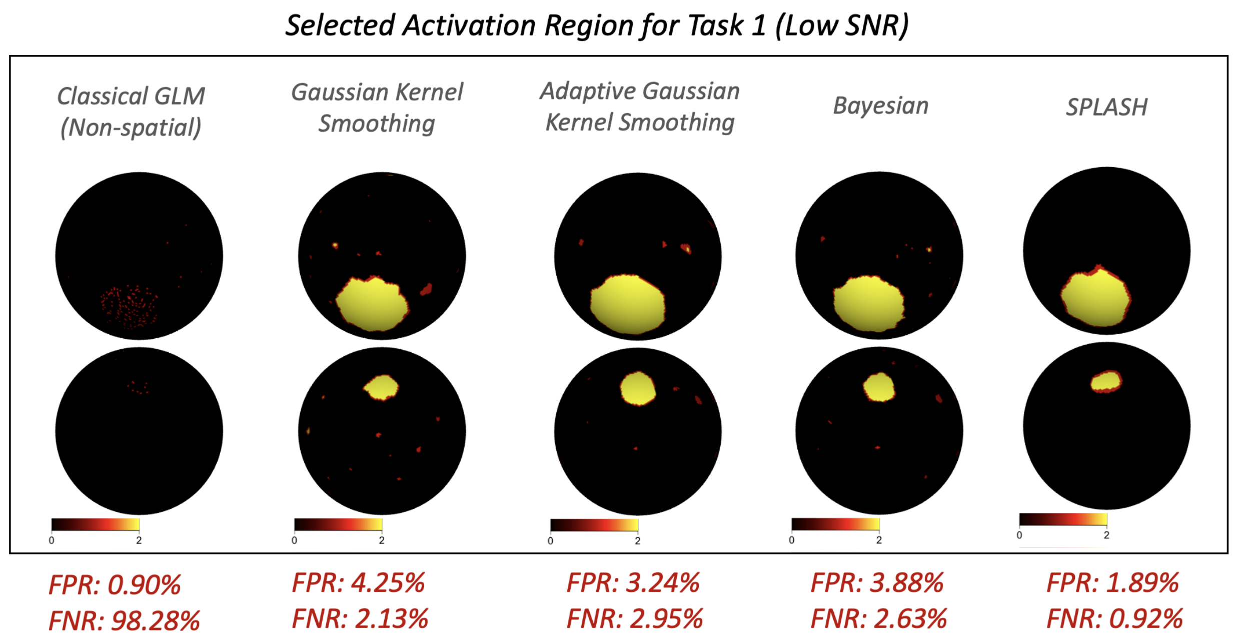

Figure S8: Detected activation for Task 1 under the low-SNR setting. False positive (FPR) and false negative (FNR) rates are shown in red.

##### 2.3.2 Task 2

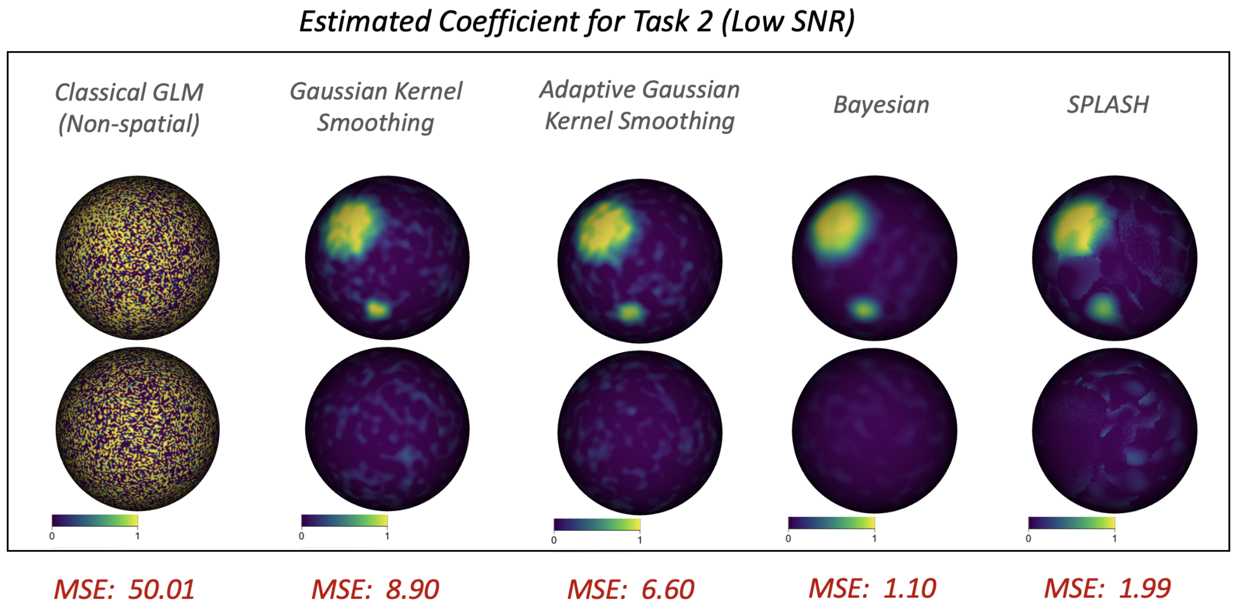

Figure S9: Estimated Task 2 coefficients under the low-SNR setting. MSE values are shown in red.

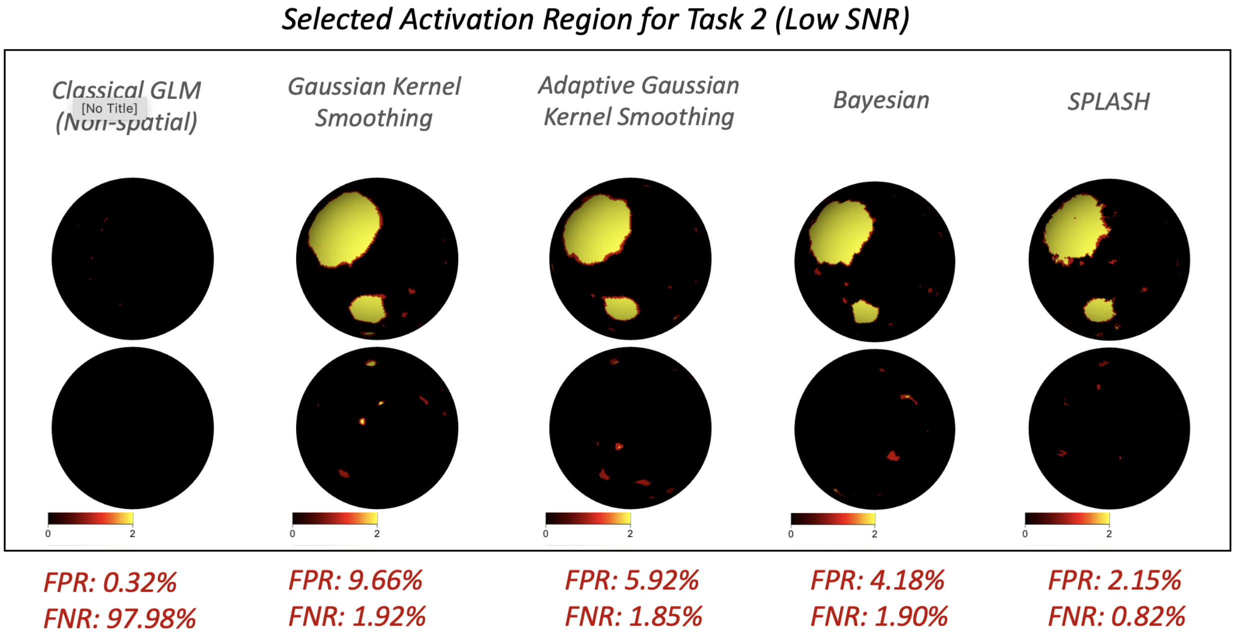

Figure S10: Detected activation for Task 2 under the low-SNR setting. False positive (FPR) and false negative (FNR) rates are shown in red.

#### 2.4 Overall Simulation Summaries

This subsection summarizes the complete set of simulation results comparing five methods across three SNR regimes (High, Moderate, and Low) and two task paradigms, complementing the results presented in Section 3.2 of the main paper. Table S1 reports estimation accuracy in terms of mean squared error (MSE) for each method and scenario. Table S2 presents inference performance, including false positive rates (FPR) and false negative rates (FNR). Finally, Table S3 examines robustness to smoothing-parameter choices by varying the Gaussian kernel bandwidth $h$ for GKS and the number of spatial spline basis functions $M$ for SPLASH.

Consistent with the visual results, the numerical summaries show that for both Task 1 and Task 2, and across all SNR settings, SPLASH achieves accurate estimation and superior activation detection. As expected, performance decreases gradually as SNR becomes lower, but SPLASH remains the most reliable method across all conditions.

We observe that SPLASH is substantially more robust to the choice of the number of spatial thin-plate splines than GKS is to its kernel bandwidth. While GKS performance is highly sensitive to the bandwidth parameter, SPLASH relies on a spline basis representation, and performance stabilizes once a sufficient number of spatial spline functions is included. Beyond this point, increasing the number of splines has minimal effect on estimation or inference quality.

Table S1: Mean squared error (MSE $\times 100$) for five methods evaluated across two tasks and three SNR conditions (low, moderate, high). Smaller values correspond to improved estimation accuracy.

| **Method** | **High SNR** | | **Moderate SNR** | | **Low SNR** | |
|---|---|---|---|---|---|---|
| | **Task 1** | **Task 2** | **Task 1** | **Task 2** | **Task 1** | **Task 2** |
| GLM | 14.86 | 12.93 | 32.47 | 34.11 | 51.62 | 50.01 |
| GKS | 4.92 | 7.72 | 3.82 | 6.35 | 6.15 | 8.90 |
| AGKS | 1.98 | 4.17 | 2.91 | 2.76 | 5.82 | 6.60 |
| Bayesian | 2.14 | 1.98 | 1.75 | 3.14 | 2.85 | 1.10 |
| SPLASH | 0.94 | 1.85 | 1.26 | 2.37 | 2.92 | 1.99 |

Table S2: False positive rate (FPR) and false negative rate (FNR) for all methods across three SNR settings (low, moderate, high) and two tasks. Lower values indicate better performance.

| **Method** | **High SNR** | | **Moderate SNR** | | **Low SNR** | |
|---|---|---|---|---|---|---|
| | **Task 1** | **Task 2** | **Task 1** | **Task 2** | **Task 1** | **Task 2** |
| **FPR (%)** | | | | | | |
| GLM | 1.62 | 3.84 | 0.78 | 3.27 | 0.90 | 0.32 |
| GKS | 9.64 | 10.65 | 10.10 | 9.29 | 4.25 | 9.66 |
| AGKS | 4.82 | 7.95 | 6.28 | 10.66 | 3.24 | 5.92 |
| Bayesian | 3.12 | 8.79 | 3.75 | 8.14 | 3.88 | 4.18 |
| SPLASH | 1.89 | 1.89 | 2.18 | 2.10 | 1.89 | 2.15 |
| **FNR (%)** | | | | | | |
| GLM | 9.21 | 32.67 | 96.17 | 67.19 | 98.28 | 97.98 |
| GKS | 0.81 | 0.93 | 1.72 | 1.00 | 2.13 | 1.92 |
| AGKS | 1.18 | 1.12 | 1.10 | 1.19 | 2.95 | 1.85 |
| Bayesian | 1.61 | 1.81 | 2.62 | 1.92 | 2.63 | 1.90 |
| SPLASH | 0.79 | 1.86 | 1.39 | 1.19 | 0.92 | 0.82 |

Table S3: Robustness to smoothing-parameter choices across three SNR levels, averaged over both tasks. For GKS, $h$ denotes the Gaussian kernel bandwidth; for SPLASH, $M$ is the number of spatial spline basis functions per parcel.

| SNR | Model (Param) | Value | FPR | FNR | MSE |
|---|---|---|---|---|---|
| High | GKS ($h$) | 0.42 | 1.78 | 10.25 | 12.81 |
| | | 0.85 | 2.75 | 8.79 | 11.43 |
| | | 1.70 | 2.28 | 6.85 | 9.72 |
| | | 2.55 | 10.15 | 0.87 | 6.32 |
| | | 3.40 | 10.92 | 0.17 | 7.50 |
| | | 4.25 | 11.52 | 0.13 | 8.91 |
| | | 5.10 | 13.92 | 0.10 | 11.69 |
| | SPLASH ($M$) | 5 | 1.89 | 0.79 | 1.46 |
| | | 10 | 1.81 | 0.79 | 1.42 |
| | | 15 | 1.83 | 0.83 | 1.49 |
| | | 20 | 1.85 | 0.80 | 1.52 |
| | | 30 | 1.90 | 0.82 | 1.57 |
| | | 40 | 1.85 | 0.79 | 1.50 |
| | | 60 | 1.85 | 0.72 | 1.38 |
| Moderate | GKS ($h$) | 0.42 | 1.88 | 10.10 | 12.37 |
| | | 0.85 | 2.63 | 8.95 | 10.82 |
| | | 1.70 | 3.45 | 7.45 | 8.54 |
| | | 2.55 | 9.72 | 1.36 | 5.09 |
| | | 3.40 | 10.85 | 0.60 | 6.47 |
| | | 4.25 | 11.41 | 0.30 | 7.72 |
| | | 5.10 | 13.90 | 0.12 | 10.18 |
| | SPLASH ($M$) | 5 | 2.19 | 1.29 | 2.42 |
| | | 10 | 2.14 | 1.24 | 2.28 |
| | | 15 | 2.23 | 1.42 | 1.87 |
| | | 20 | 2.04 | 1.20 | 2.06 |
| | | 30 | 2.97 | 1.35 | 2.62 |
| | | 40 | 3.02 | 1.40 | 2.75 |
| | | 60 | 2.10 | 1.96 | 2.48 |
| Low | GKS ($h$) | 0.42 | 1.53 | 10.20 | 13.42 |
| | | 0.85 | 2.11 | 9.15 | 11.96 |
| | | 1.70 | 3.01 | 7.85 | 9.84 |
| | | 2.55 | 6.96 | 2.03 | 7.53 |
| | | 3.40 | 8.44 | 1.10 | 8.62 |
| | | 4.25 | 9.57 | 0.55 | 9.87 |
| | | 5.10 | 11.82 | 0.25 | 12.44 |
| | SPLASH ($M$) | 5 | 2.89 | 0.87 | 2.18 |
| | | 10 | 2.01 | 0.91 | 1.96 |
| | | 15 | 2.84 | 1.01 | 2.54 |
| | | 20 | 2.90 | 1.72 | 2.88 |
| | | 30 | 2.17 | 0.99 | 2.12 |
| | | 40 | 2.76 | 0.89 | 2.33 |
| | | 60 | 2.65 | 0.92 | 2.25 |

### 3 Additional Application Results

This section presents subject-level and group-level results for the Left Foot, Right Foot, Left Hand, and Right Hand tasks under the Schaefer-100 parcellation. Corresponding analyses for the remaining motor tasks discussed in Section 4.1 of the main paper, as well as SPLASH results under alternative parcellations, are provided in Figures S11–S22.

We observe that even at the single-subject level, SPLASH produces clearer and more spatially coherent activation maps when paired with selective inference. At the group level, SPLASH yields activation patterns that show stronger correspondence with the known motor homunculus. Moreover, SPLASH is not overly dependent on the choice of parcellation: across all motor tasks, both the detected activation regions and the estimated activation coefficients remain highly consistent across different parcellation schemes.

#### 3.1 Single-Subject Analysis

Even if it's

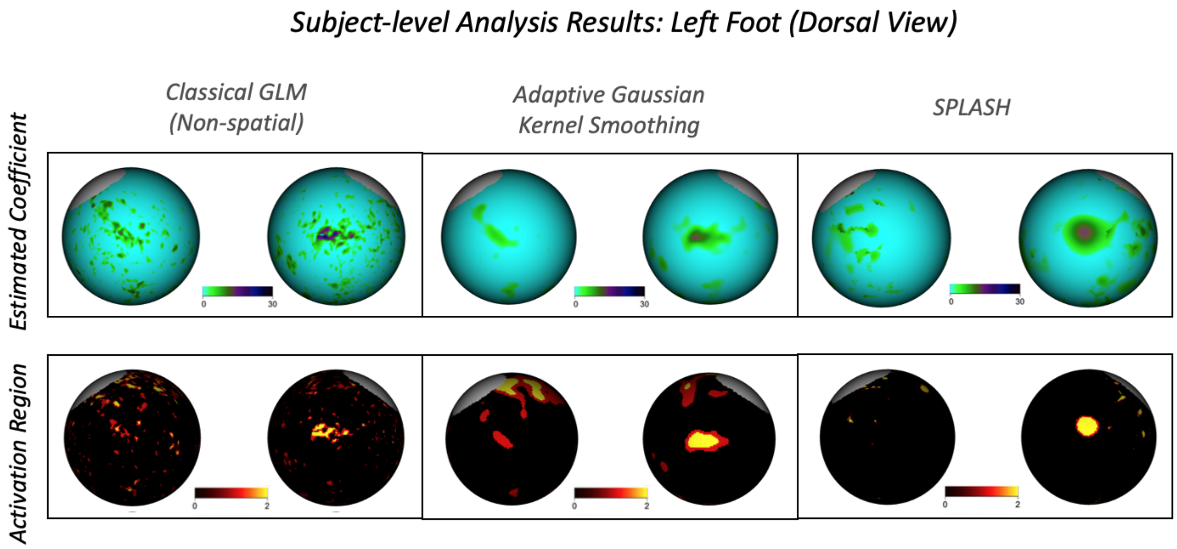

Figure S11: Top row: Estimated coefficients for the left-foot task for single subject. Bottom row: Corresponding detected activation regions (yellow: $p < 0.01$, red: $p < 0.05$).

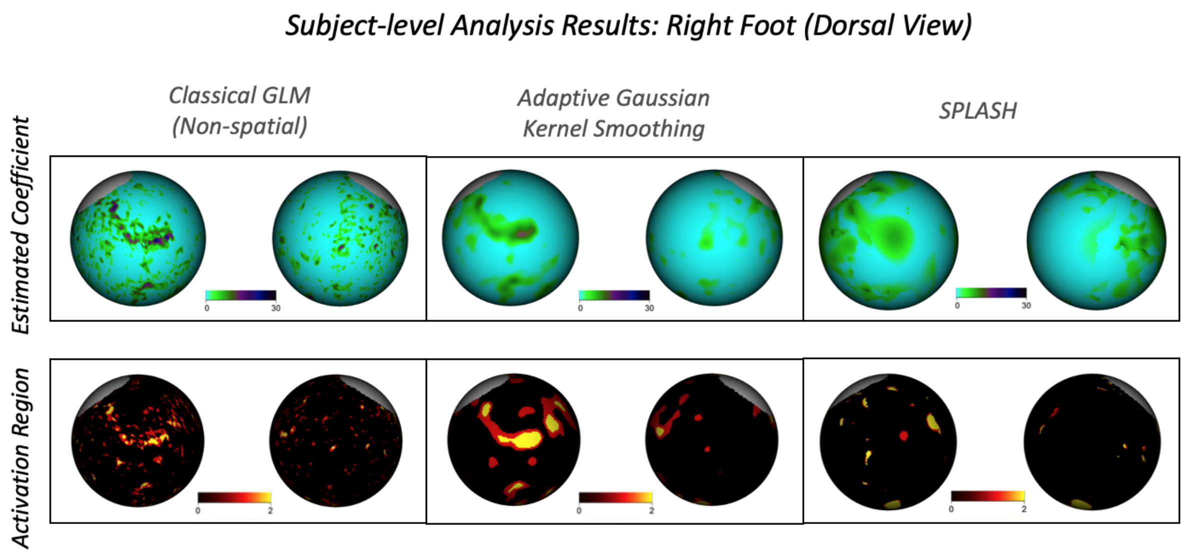

Figure S12: Top row: Estimated coefficients for the right-foot task for single subject. Bottom row: Corresponding detected activation regions (yellow: $p < 0.01$, red: $p < 0.05$).

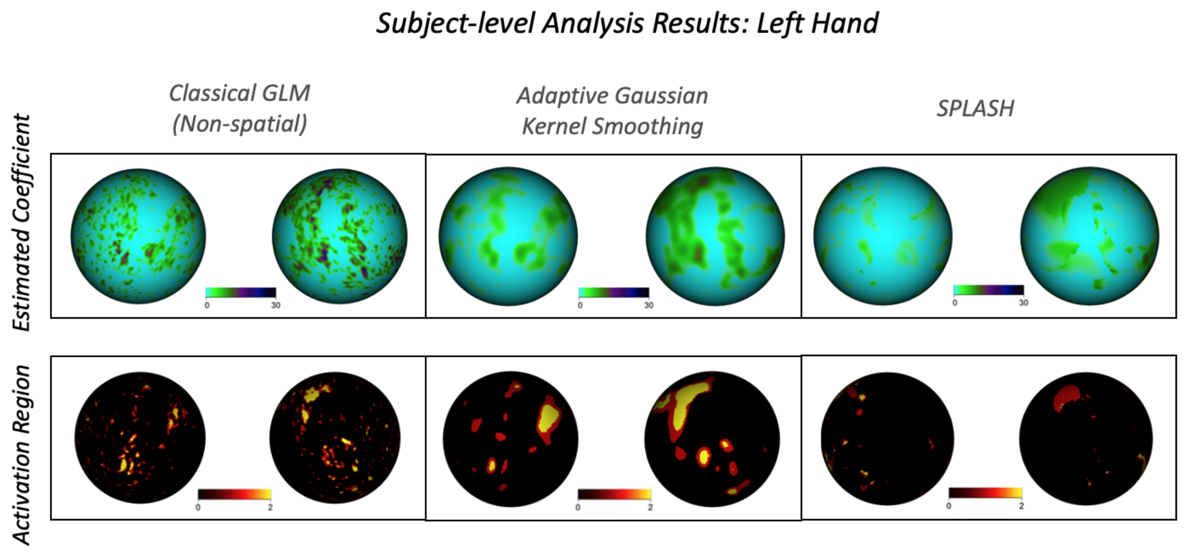

Figure S13: Top row: Estimated coefficients for the left-hand task for single subject. Bottom row: Corresponding detected activation regions (yellow: $p < 0.01$, red: $p < 0.05$).

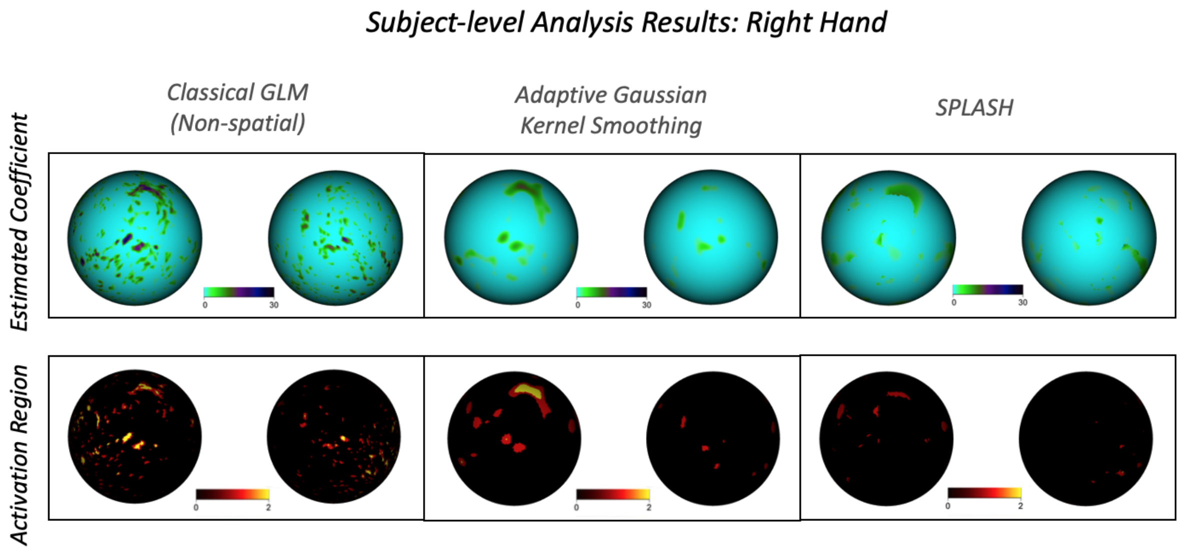

Figure S14: Top row: Estimated coefficients for the right-hand task for single subject. Bottom row: Corresponding detected activation regions (yellow: $p < 0.01$, red: $p < 0.05$).

#### 3.2 Multi-Subject Group Analysis

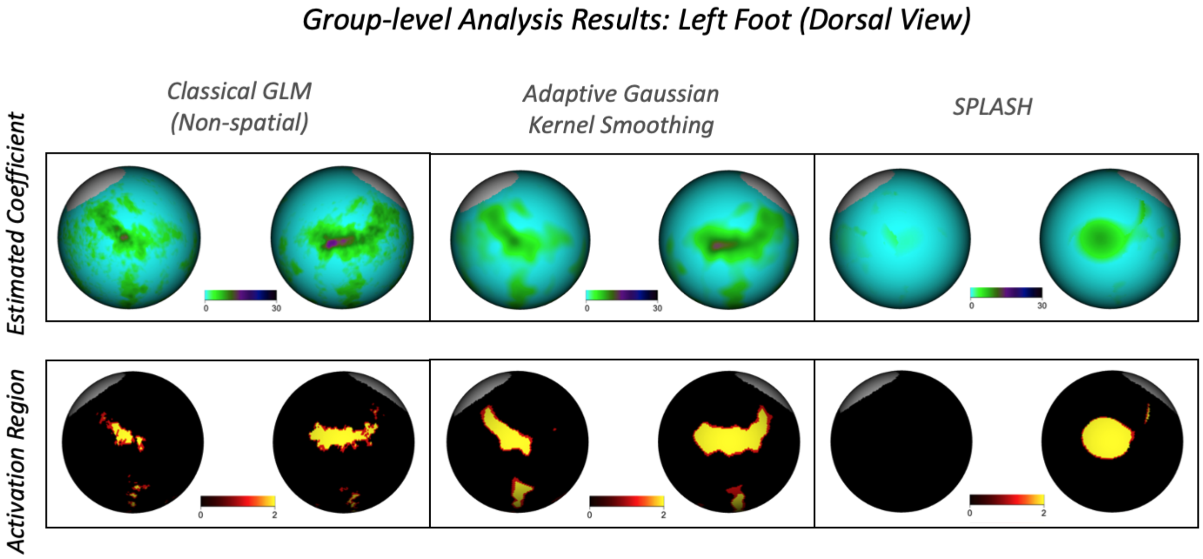

Figure S15: Top row: Estimated coefficients for the left-foot task for 416 subjects. Bottom row: Corresponding detected activation regions (yellow: $p < 0.01$, red: $p < 0.05$).

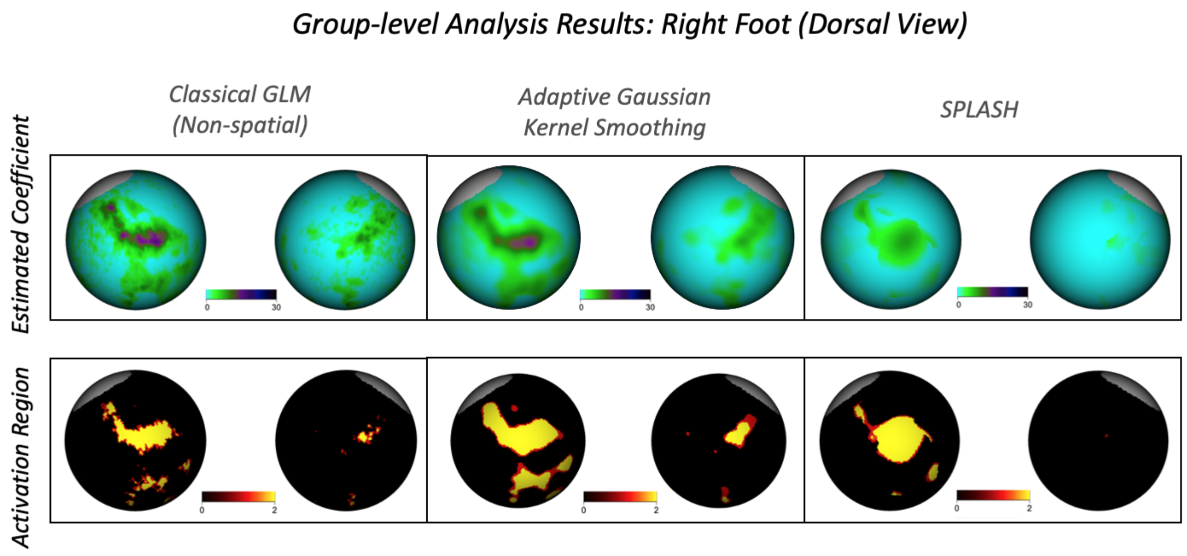

Figure S16: Top row: Estimated coefficients for the right-foot task for 416 subjects. Bottom row: Corresponding detected activation regions (yellow: $p < 0.01$, red: $p < 0.05$).

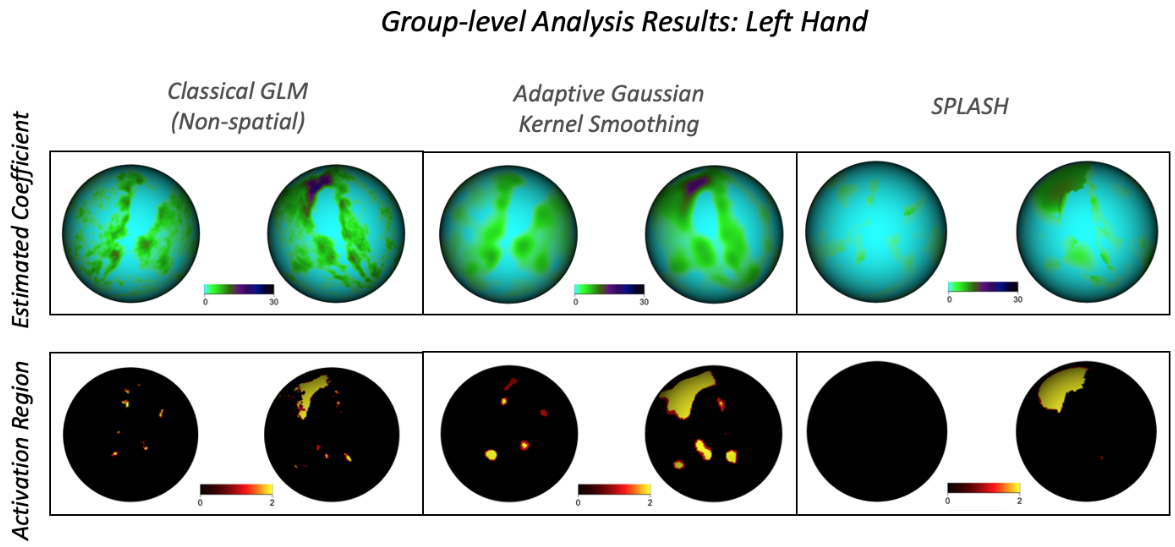

Figure S17: Top row: Estimated coefficients for the left-hand task for 416 subjects. Bottom row: Corresponding detected activation regions (yellow: $p < 0.01$, red: $p < 0.05$).

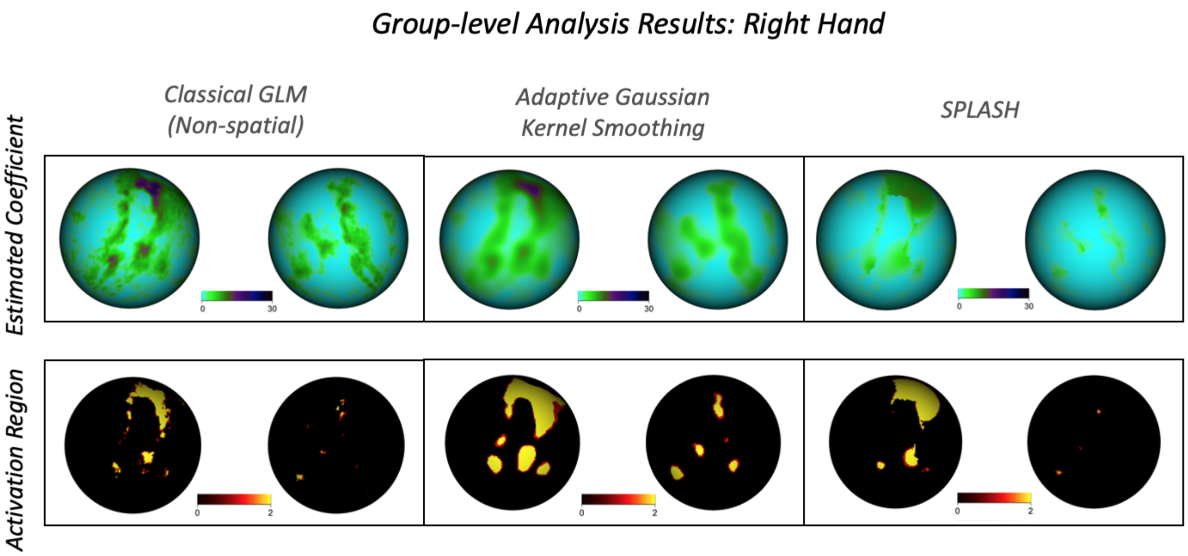

Figure S18: Top row: Estimated coefficients for the right-hand task for 416 subjects. Bottom row: Corresponding detected activation regions (yellow: $p < 0.01$, red: $p < 0.05$).

#### 3.3 Robustness to Parcellation

This subsection presents robustness-to-parcellation visualizations for the four remaining motor tasks (Left Foot, Right Foot, Left Hand, and Right Hand) under the three parcellation schemes described in Section 3.2.3 of the main paper for the Tongue task.

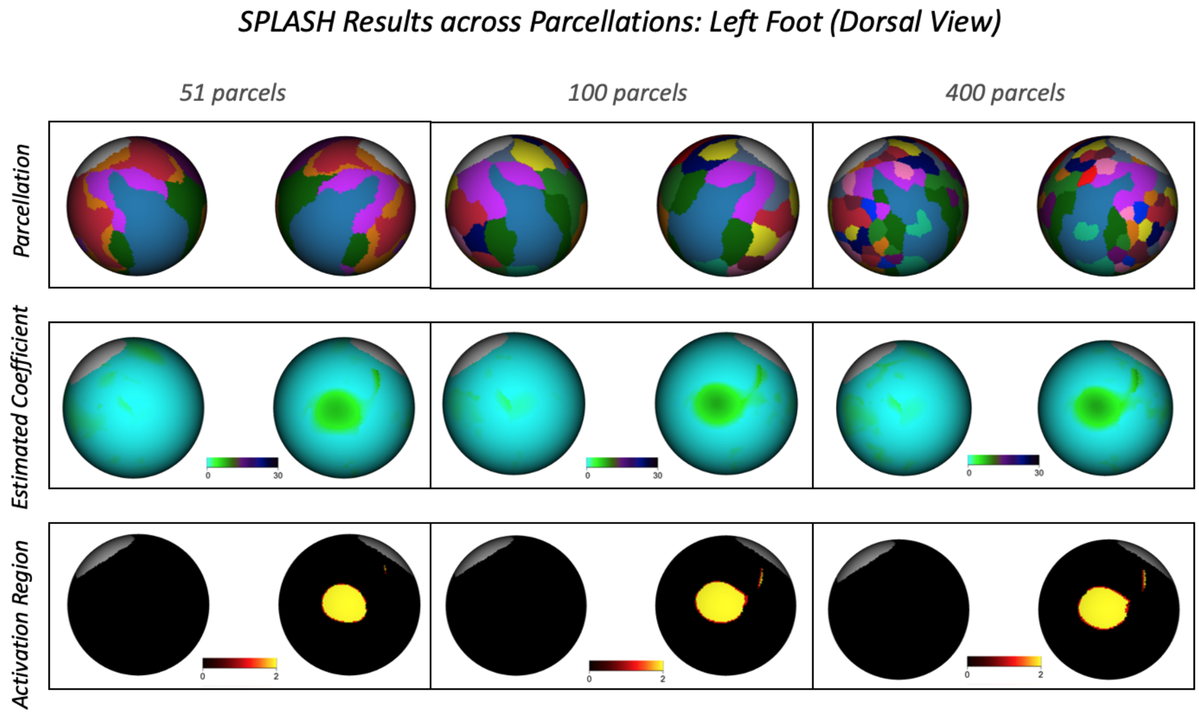

Figure S19: Top row: cortical parcellations used in the robustness analysis. Middle row: SPLASH-estimated activation coefficients for each corresponding parcellation for left-foot task. Bottom row: detected activation regions (yellow: $p < 0.01$, red: $p < 0.05$).

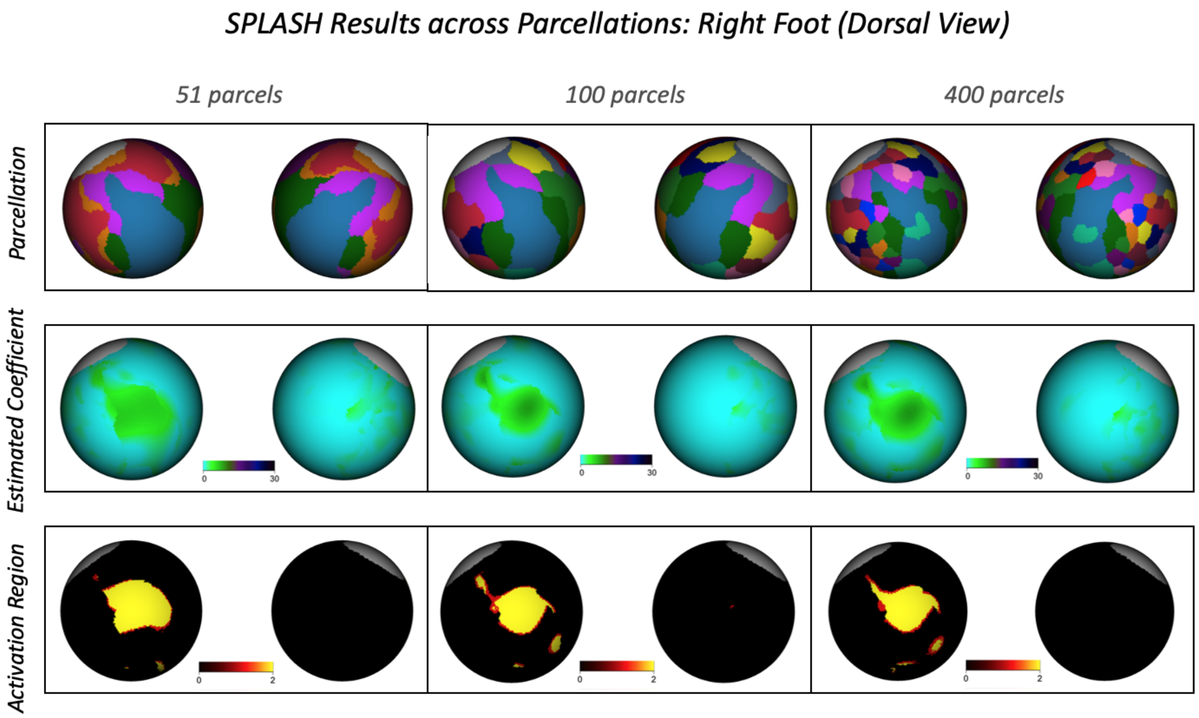

Figure S20: Top row: cortical parcellations used in the robustness analysis. Middle row: SPLASH-estimated activation coefficients for each corresponding parcellation for right-foot task. Bottom row: detected activation regions (yellow: $p < 0.01$, red: $p < 0.05$).

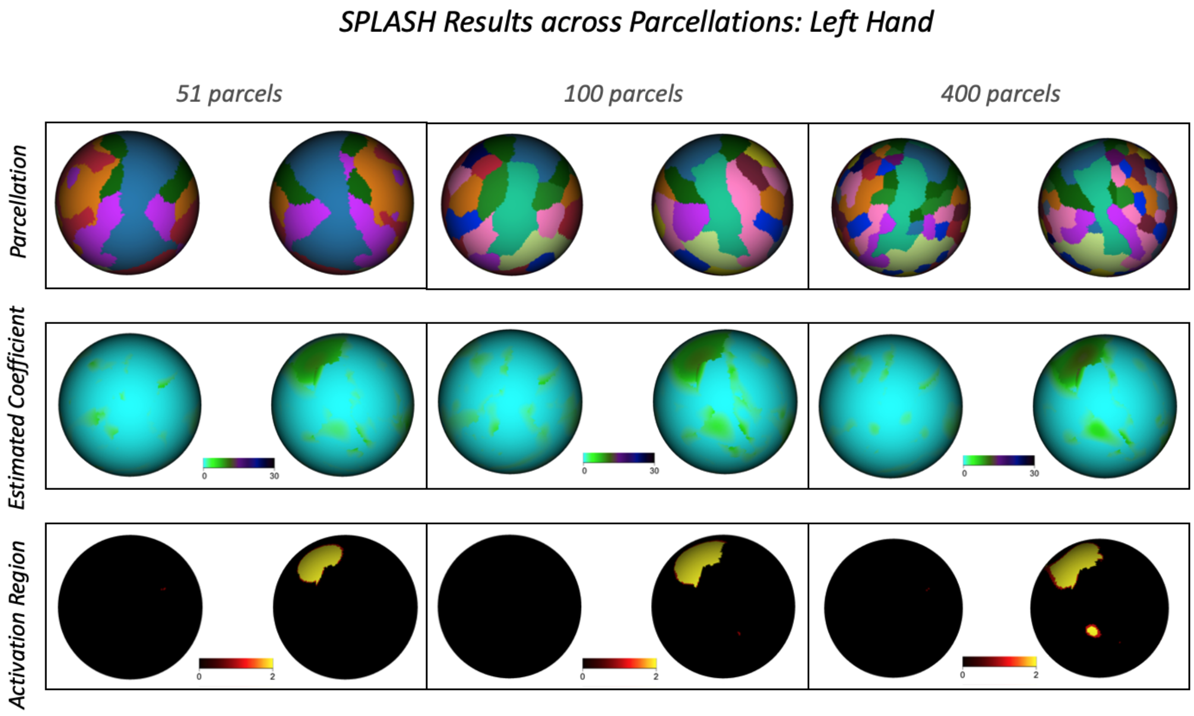

Figure S21: Top row: cortical parcellations used in the robustness analysis. Middle row: SPLASH-estimated activation coefficients for each corresponding parcellation for left-hand task. Bottom row: detected activation regions (yellow: $p < 0.01$, red: $p < 0.05$).

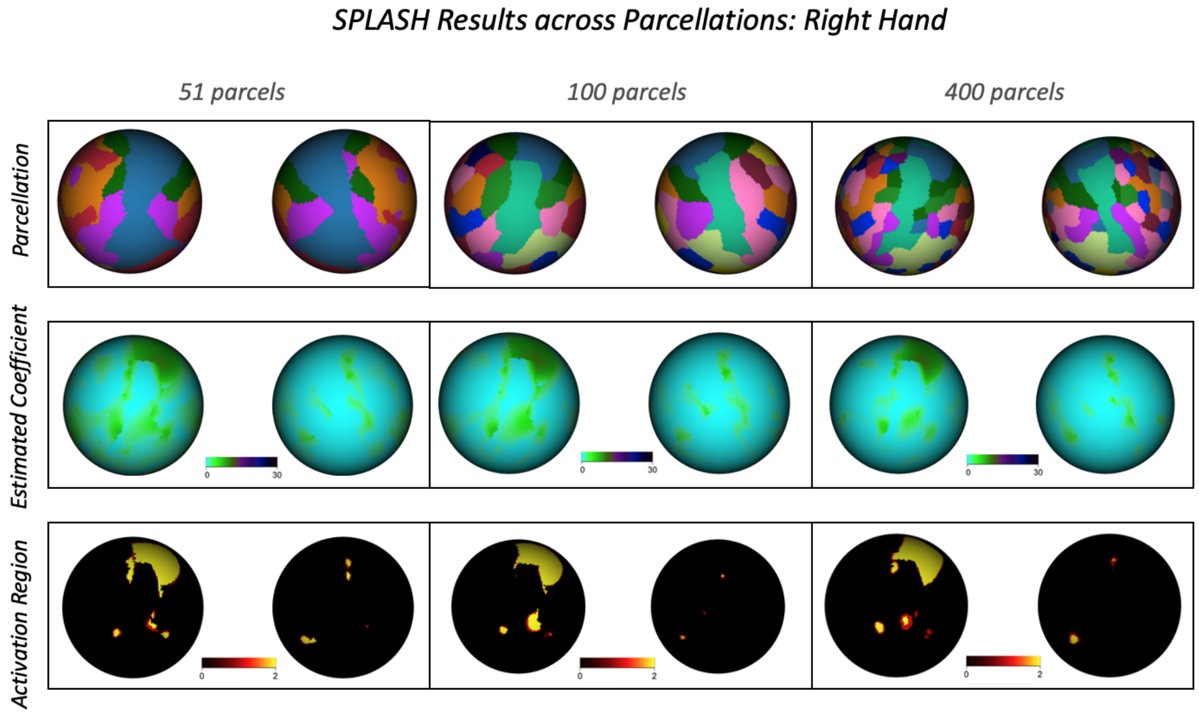

Figure S22: Top row: cortical parcellations used in the robustness analysis. Middle row: SPLASH-estimated activation coefficients for each corresponding parcellation for right-hand task. Bottom row: detected activation regions (yellow: $p < 0.01$, red: $p < 0.05$).
